# Pilotins are mobile T3SS components involved in assembly and substrate specificity of the bacterial type III secretion system

**DOI:** 10.1101/2022.02.14.480308

**Authors:** Stephan Wimmi, Moritz Fleck, Carlos Helbig, Corentin Brianceau, Katja Langenfeld, Witold G. Szymanski, Georgia Angelidou, Timo Glatter, Andreas Diepold

## Abstract

In animal pathogens, assembly of the type III secretion system injectisome requires the presence of so-called pilotins, small lipoproteins that assist the formation of the secretin ring in the outer membrane. Using a combination of functional assays, interaction studies, proteomics, and live-cell microscopy, we determined the contribution of the pilotin to the assembly, function, and substrate selectivity of the T3SS and identified potential new downstream roles of pilotin proteins. In absence of its pilotin SctG, *Yersinia enterocolitica* forms few, largely polar injectisome sorting platforms and needles. In line, most export apparatus subcomplexes are mobile in these strains, suggesting the absence of fully assembled injectisomes. Remarkably, while absence of the pilotin all but prevents export of early T3SS substrates, such as the needle subunits, it has little effect on secretion of late T3SS substrates, including the virulence effectors. We found that pilotins transiently interact with other injectisome components such as the secretin in the outer membrane, but mostly form transient mobile clusters in the bacterial membrane, which do not colocalize with assembled injectisomes. Together, these findings provide a new view on the role of pilotins during and after assembly of type III secretion injectisomes.

## Introduction

Bacteria that live in contact to eukaryotic cells greatly benefit from being able to manipulate those cells. The bacterial type III secretion system (T3SS) allows such manipulation by injecting effector proteins directly from the bacterial cytosol into the host cell cytoplasm [1–5]. Although the T3SS is also used by symbionts and commensals, it is best known for its essential role in infections of important human pathogens such as *Salmonella, Shigella* and *Yersinia*. In these pathogens, the T3SS machinery, also called injectisome^1^, is often assembled upon entry into the host organism. Once the basal body (membrane rings and export apparatus, Fig. 1a) and the cytosolic components are assembled (reviewed in [6]), the T3SS starts to secrete its own distal components. Amongst those so-called early secretion substrates are the subunits of the needle and proteins required for its correct assembly, as well as early regulatory proteins. When the required needle length is reached [7,8], middle secretion substrates – the translocon proteins forming the needle tip and two hydrophobic interactors that build a pore in the host membrane – are secreted. At this point, the injectisome is ready for secretion of the late secretion substrates, the virulence effectors, which is initiated by host cell contact.

**Fig. 1.**
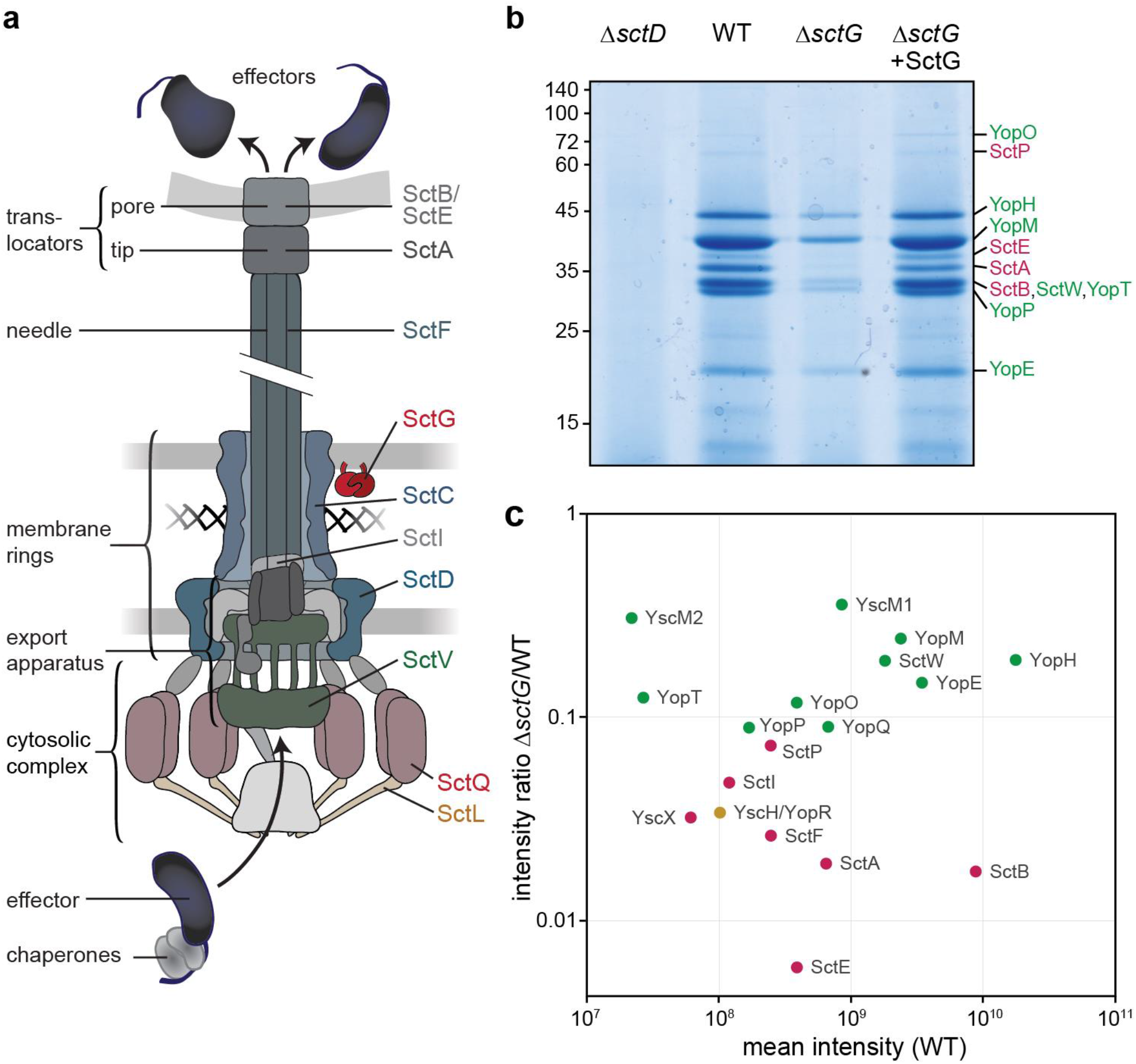
SctG is required for the secretion of early export substrates, but not of effectors. **a**) Schematic illustration of the injectisome (adapted from [6]). Main substructures indicated on the left; components analyzed in this study, including the pilotin SctG (for which a possible location is indicated, see main text) and the secretin SctC, are indicated on the right and shown in color. **b**) Secretion assay showing the export of native T3SS substrates into the culture supernatant in the listed strains in rich medium (*n*=3). Left, molecular weight in kDa; right, assignment of exported proteins. **c**) Label-free mass spectrometry quantification of secreted proteins in *ΔsctG* and wild-type (WT) in rich medium, intensity ratio plotted against mean intensity (*n*=3). In b) and c), early/middle T3SS export substrates are marked in magenta; late substrates (effectors) are marked in green; substrate with unknown time point of export marked in yellow. See Fig. S1 for comparison between rich and minimal medium.

Injectisomes are assembled at the bacterial surface according to the needs of the bacteria in a given situation. While in some cases, one to few injectisomes are sufficient (such as for the intracellular *Salmonella* SPI-2 T3SS [9]), bacteria that need to actively make contact with the host cells usually build more injectisomes [10,11]. The distribution of injectisomes across the bacterial surface is non-random [12–14], but it is currently unclear what determines where new injectisomes are built.

A prime candidate to establish the site of assembly for new injectisomes is the pilotin, a lipoprotein anchored in the inner leaflet of the outer membrane (OM). Pilotins are required for the proper integration and oligomerization of the secretin protein, which forms a large channel in the OM in the T3SS of animal pathogens, but also type II secretion systems (T2SS) and type IV pili (T4P) [15–24]. Since the secretin ring is one of the nucleation structures of the injectisome [6,25] and anchors the T3SS at its final position in the peptidoglycan mesh [26] (Fig. 1a), the localization of the pilotin may determine the place of secretin ring formation and in consequence the distribution of injectisomes.

Pilotins and their cognate secretin proteins in the T2SS and T3SS have been purified and structurally analyzed in 1:1 complexes [17,27,28]. As a lipoprotein, the pilotin also interacts with the localization of lipoproteins (Lol) protein export machinery [28,29]. Based on these properties, different models for the export of the pilotin itself and its role in secretin ring assembly have been proposed [27,28]. Whether the pilotin stays attached to the secretin after secretin ring formation in the OM and the subsequent assembly of the injectisome is debated and might differ between organisms. For T2SS secretins, such attachment was initially proposed, but later rejected [17,30]; recent analyses of purified secretin rings found pilotins attached stoichiometrically outside the main ring [31,32]. *In situ* structures of the T3SS showed electron densities around the secretin, possibly corresponding to pilotin proteins, in *Shigella flexneri* [33,34], but not in *Salmonella enterica* [35–37]. In spite of the importance of pilotins for the localization of the central secretin ring structure, protein secretion by the T3SS was found to be strongly impaired, but not completely prevented in absence of the pilotin in different organisms [15,19,24]. Likewise, the number of needles formed was strongly reduced in a pilotin mutant in *P. aeruginosa* [24]. The impact on secretion varied between different bacteria and substrates, and the structural and mechanistic basis for this phenotype is currently unclear. Pilotins have a conserved unique genetic location within the virulence plasmids or genetic islands encoding the T3SS. While most other structural and regulatory components of the injectisome are encoded in operons under the control of a main transcriptional regulator (VirF in *Y. enterocolitica*, called LcrF in other *Yersinia* species), pilotins are located upstream of the transcriptional regulator itself, indicating a specific role in the T3SS with a potentially distinct expression pattern.

Despite their central role, many key characteristics of pilotins are poorly understood or debated: Why are pilotins encoded differently from all other T3SS components? When and where are they active and does this determine the assembly and localization of injectisomes? Do pilotins remain part of the injectisome after the assembly and do they adapt a functional role in type III secretion at this point?

To shed light on these questions and the role of pilotins in assembly and function of the T3SS, we studied the role of the *Yersinia enterocolitica* pilotin SctG^2^ with functional assays, fluorescence microscopy, proteomics, as well as protein interaction and dynamics studies. We found that, unexpectedly, SctG interacts with the secretin SctC only temporarily, and mostly localizes in dynamic patches that are dispersed throughout the bacterial membrane. In absence of the pilotin, only a small subset of bacteria assemble single, mostly polar needles which are not stably attached to fully assembled injectisomes, as indicated by the lack of colocalization with the cytosolic sorting platform component SctL. Most notably, presence of the pilotin not only influenced injectisome assembly and overall secretion levels, but specifically altered the secretion pattern of the T3SS. While effector proteins and other late substrates were efficiently secreted in the absence of the pilotin, the export of early substrates, such as the needle components, was greatly impaired. Based on specific interaction studies, we provide possible explanations for this unique phenotype and an extended model for the function of the pilotin.

## Results

### Deletion of the pilotin specifically reduces secretion of early export substrates

To define the overall role of the pilotin SctG in type III secretion, we tested the effects of a complete deletion of the *sctG* gene on the virulence plasmid in *Y. enterocolitica*. We found that absence of SctG distinctly altered the secretion pattern. While secretion of effector proteins was reduced, but still clearly present, early secretion substrates such as the needle subunit SctF, the ruler protein SctP and translocators SctA, B, E were much more severely affected in secretion or not secreted at all. *In trans* expression of SctG complemented the deletion (Fig. 1b, Fig. S1a).

In order to determine the role of SctG in secretion more precisely, we analyzed the secretome of the wild-type and *ΔsctG* strains by label-free quantitative mass spectrometry. The results confirm the secretion assay and show that T3SS substrates cluster according to their substrate class, rather than their overall secretion level (Fig. 1c, Table 1, Fig. S1a): The late substrates, effector proteins YopH, O, P, E, M, T as well as the gatekeeper SctW (YopN) and the negative regulators YscM1/2 (marked green in Fig. 1bc), were exported efficiently in absence of SctG. In clear contrast, export of all early and middle substrates of the T3SS (marked magenta in Fig. 1bc) was strongly reduced in *ΔsctG*. These T3SS substrates build up the needle (SctF) and the inner rod (SctI) connecting it to the export apparatus, control needle length (SctP) and form its tip structure (SctA) as well as a pore in the host membrane (SctB, E). YscX is an early T3SS substrate specific to the Ysc subgroup of T3SS [38,39], which is required for the export of any other protein by the T3SS of the Ysc subgroup [40,41]. Notably, secretion of the late substrates is fully T3SS-dependent, as no secretion was observed in a *ΔsctD* strain (Fig. 1b). In minimal medium, which allows for a more quantitative analysis by proteomics approaches, overall secretion levels were reduced, but an even more pronounced selective export of late substrates could be observed (Table 1, Fig. S1ab).

**Table 1.** Influence of SctG on T3SS secretion substrates. Label-free quantitative mass spectrometry of known T3SS substrates in the culture supernatant of different strains, as indicated. Top, measurement in rich medium (BHI); bottom, in minimal medium (see material and methods for details). Magenta shade, early and middle substrates; green shade, late substrates; yellow shade, unknown. # pept., number of total detected peptides. Light grey font, imputed values (no peptides detected in at least one sample). Hits sorted by ratio of *ΔsctG*/wild-type (WT). *n*=3, individual measurements listed in Table S3.

| *Rich medium* | #<br>pept. | Intensity<br>ratio ΔsctG /<br>WT | Mean intensity | | |
| --- | --- | --- | --- | --- | --- |
| T3SS substrate | | | WT | ΔsctG | ΔsctD |
| Translocator SctE (YopB) | 29 | 0.0059 | 3.89E+08 | 2.29E+06 | 3.27E+05 |
| Translocator SctB (YopD) | 73 | 0.0174 | 8.87E+09 | 1.54E+08 | 2.76E+06 |
| Tip protein SctA (LcrV) | 41 | 0.0190 | 6.48E+08 | 1.23E+07 | 7.97E+04 |
| Needle component SctF | 8 | 0.0261 | 2.47E+08 | 6.45E+06 | 1.61E+05 |
| Early substrate / regulator YscX | 9 | 0.0321 | 6.09E+07 | 1.95E+06 | 3.06E+05 |
| Secreted protein YscH/YopR | 8 | 0.0339 | 1.01E+08 | 3.43E+06 | 5.17E+04 |
| Rod / washer protein SctI | 8 | 0.0476 | 1.19E+08 | 5.66E+06 | 5.20E+04 |
| Needle length regul. / ruler SctP | 33 | 0.0724 | 2.46E+08 | 1.78E+07 | 7.78E+05 |
| Effector YopP | 41 | 0.0886 | 1.68E+08 | 1.49E+07 | 7.50E+04 |
| Effector YopQ | 24 | 0.0895 | 6.70E+08 | 6.00E+07 | 8.07E+05 |
| Effector YopO | 71 | 0.1178 | 3.87E+08 | 4.55E+07 | 1.21E+05 |
| Effector YopT | 13 | 0.1246 | 2.64E+07 | 3.29E+06 | 4.27E+04 |
| Effector YopE | 25 | 0.1473 | 3.47E+09 | 5.10E+08 | 7.83E+05 |
| Gatekeeper SctW (YopN) | 19 | 0.1890 | 1.81E+09 | 3.42E+08 | 1.06E+06 |
| Effector YopH | 122 | 0.1906 | 1.78E+10 | 3.39E+09 | 3.17E+06 |
| Effector YopM | 8 | 0.2428 | 2.39E+09 | 5.81E+08 | 1.46E+06 |
| Regulator YscM2 | 9 | 0.3061 | 2.16E+07 | 6.61E+06 | 5.69E+04 |
| Regulator YscM1 | 21 | 0.3562 | 8.54E+08 | 3.04E+08 | 7.61E+05 |

| *Minimal medium* | #<br>pept. | Intensity<br>ratio ΔsctG /<br>WT | Mean intensity | | |
| --- | --- | --- | --- | --- | --- |
| T3SS substrate | | | WT | ΔsctG | ΔsctC |
| Needle component SctF | 7 | 0.0547 | 3.95E+09 | 2.16E+08 | 1.65E+07 |
| Early substrate / regulator YscX | 4 | 0.1366 | 3.55E+08 | 4.85E+07 | 8.80E+06 |
| Tip protein SctA (LcrV) | 24 | 0.1580 | 8.72E+09 | 1.38E+09 | 1.07E+08 |
| Rod / washer protein SctI | 3 | 0.1621 | 3.56E+08 | 5.78E+07 | 2.07E+07 |
| Needle length regul. / ruler SctP | 23 | 0.2359 | 1.53E+09 | 3.60E+08 | 4.84E+07 |
| Translocator SctE (YopB) | 19 | 0.3213 | 2.73E+09 | 8.77E+08 | 3.44E+08 |
| Translocator SctB (YopD) | 46 | 0.4502 | 3.26E+10 | 1.47E+10 | 9.06E+08 |
| Secreted protein YscH/YopR | 4 | 0.5604 | 2.90E+08 | 1.62E+08 | 1.15E+07 |
| Effector YopQ | 16 | 0.6357 | 1.90E+09 | 1.21E+09 | 6.51E+06 |
| Effector YopO | 37 | 0.9833 | 1.72E+09 | 1.69E+09 | 6.58E+07 |
| Effector YopH | 64 | 1.1095 | 3.11E+10 | 3.45E+10 | 1.23E+09 |
| Effector YopE | 10 | 1.5746 | 1.16E+10 | 1.82E+10 | 2.67E+09 |
| Effector YopP | 25 | 2.1810 | 7.21E+08 | 1.57E+09 | 5.24E+06 |
| Gatekeeper SctW (YopN) | 17 | 2.5811 | 3.65E+09 | 9.42E+09 | 3.18E+07 |
| Effector YopM | 6 | 2.6303 | 4.81E+09 | 1.26E+10 | 7.99E+07 |
| Effector YopT | 7 | 4.1605 | 1.15E+07 | 4.77E+07 | 1.28E+07 |
| Regulator YscM1 | 12 | 5.5968 | 4.84E+08 | 2.71E+09 | 9.81E+06 |
| Regulator YscM2 | 7 | 10.2833 | 4.52E+07 | 4.65E+08 | 5.43E+06 |

Most pilotins, including SctG, are expressed in an operon with the main transcription factor of the T3SS, VirF, which is encoded downstream of *sctG* in the same operon and whose expression is controlled by an RNA thermometer structure located between the two genes [42,43]. We therefore tested the influence of the *sctG* deletion on VirF expression levels. In *ΔsctG*, the amount of the main transcriptional activator VirF was reduced to about one third. Consequently, most other T3SS components were also present at lower concentrations (Table S2). To test if the observed phenotype was caused by this impaired VirF expression in the absence of *sctG*, we expressed VirF from plasmid in a *ΔsctG* strain. Indeed, this slightly increased the secretion of effectors, but not of early substrates (Fig. S1c), indicating that the specific loss of secretion of these early substrates is not caused by altered VirF expression in absence of *sctG*. Importantly, these results highlight that the selective export observed in the absence of the pilotin (Fig. 1bc, Table 1) is not due to differential expression of the respective export cargo (Table S2).

### The pilotin is essential for stable localization of the export apparatus and the cytosolic T3SS components, but few needles can form in its absence

We next investigated the specific role of the pilotin in the assembly of the injectisome, which may provide a clue to its function. To this aim, we deleted *sctG* in strains expressing fluorescently labeled variants of the secretin (SctC-mCherry) [25], the large export apparatus component (SctV-EGFP) [26], the cytosolic sorting platform component SctL (mCherry-SctL) [44], and a strain expressing SctF_S5C_, a needle protein variant that can be labeled by maleimide dyes [45]. Except for SctF_S5C_, which is expressed from a plasmid, all other labeled proteins are expressed from their native genetic environment. As expected, fluorescence microscopy of the respective bacteria showed an important role of SctG in the assembly of the secretin ring. Far from its usual assembly in >10 mostly lateral foci in the wild-type background, SctC mostly localized in large, often polar clusters in the absence of the pilotin (Fig. 2a, left, Dataset S1). We found that these SctC foci generally do not correspond to fully assembled injectisomes: The cytosolic component SctL formed very few foci in the absence of SctG (Fig. 2, center; Dataset S1), and SctV foci were motile in the membrane in the absence of SctG, like in the absence of the secretin SctC (Fig. 2b, Fig. S2a) [26]. Nevertheless, a small subset of bacteria lacking SctG did display needles (Fig. 2, right; Dataset S1). These needles, however, did not colocalize with the sorting platform protein SctL (Fig. S2b). *In trans* complementation of SctG restored wild-type localization of all tested components (Fig. S3). Notably, levels of SctG significantly lower than in the wild-type were already sufficient for complementation of secretion and injectisome assembly (Fig. S3).

**Fig. 2.**
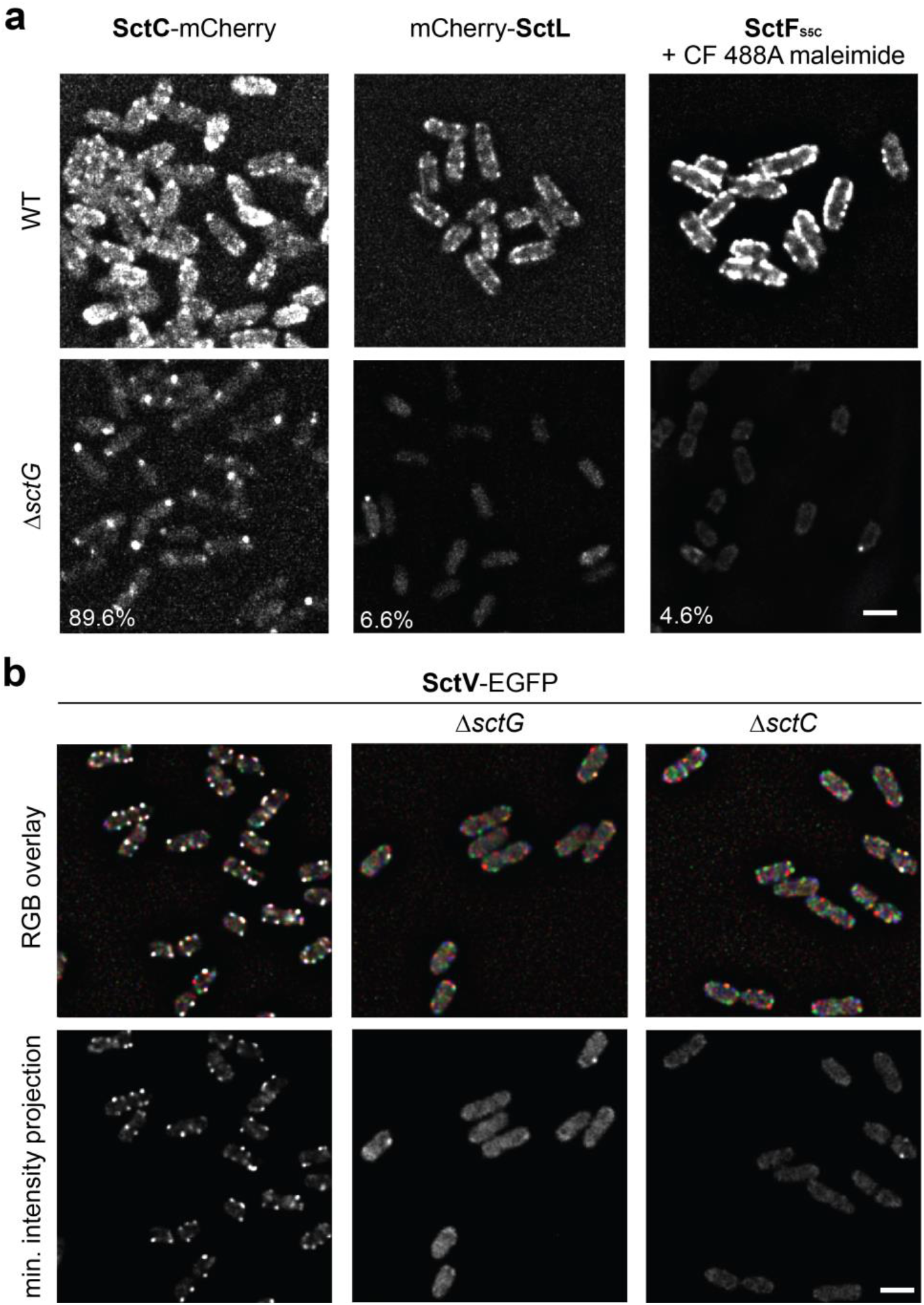
SctG is required for normal localization of SctC and stable assembly of the export apparatus and the sorting platform component SctL, but not fully essential for the formation of needles. **a**) Fluorescence micrographs of *Y. enterocolitica* expressing the proteins indicated from their native location on the virulence plasmid (SctC-mCherry and mCherry-SctL) or *in trans* (SctF_S5C_, stained with CF 488A maleimide stain) in an effector-less strain background. Top row displays native localization in wildtype (WT) background, bottom row localization in respective *ΔsctG* strain background. Scale bar, 2 μm. The fraction of bacteria with foci is listed for the respective *ΔsctG* strains. See Dataset S1 for details. *n*=3-6. **b**) Red/green/blue (RGB) overlay and minimum intensity projection showing the movement of SctV-EGFP over time in the indicated effector-less strain backgrounds in live *Y. enterocolitica*. Red/green/blue channel: micrograph at t=0/15/30 s; white foci indicate stable protein localization, whereas colors indicate movement of foci over time. Minimum projection: micrographs at 0/15/30/45/60/75/90 s; foci indicate stable foci over this time period. *n*=3.

The results above indicate that SctG is essential for proper assembly and anchoring of all major subcomplexes of the injectisome, but that a subset of cells form mostly polar needles in its absence. We therefore took a closer look at the interactions, expression kinetics and localization of SctG, to figure out how it conveys its function.

### Proteome analysis and specific interactions of the pilotin to T3SS components and proteins not directly linked to the T3SS

The finding that SctG amounts below the wild-type level are sufficient for its function in injectisome assembly (Fig. S3) prompted us to consider additional roles of the pilotin. To identify such roles in an unbiased way, we compared the full *Y. enterocolitica* proteome in presence and absence of SctG under non-secreting conditions. Absence of SctG had little impact beyond the T3SS, with the notable exception of the OM protein OmpW, whose amount was reduced by a factor of eight (Table S2).

To unravel the expression kinetics, as well as the subcellular localization of SctG, we introduced a *sctG-sfGFP* fusion in the native *sctG* locus on the *Y. enterocolitica* virulence plasmid by homologous recombination [46]. The SctG-sfGFP fusion protein was stable and functional (Fig. S4). The expression of SctG itself, VirF and, in consequence, other T3SS components was moderately increased (1.5-2-fold) in the strain. Additionally, few non-T3SS proteins were affected by the fusions (Table S4).

In an attempt to better understand the function of the pilotin, both with respect to the T3SS and other pathways, we then screened for interaction partners by co-immunoprecipitation with SctG-sfGFP, analyzed by label-free mass spectrometry. Confirming the specificity of the co-immunoprecipitation experiment, SctG interacted with the secretin SctC, but also with other T3SS components, most prominently the effector YopM and the large export apparatus protein SctV (Fig. 3). Most other T3SS components were enriched to a lower degree (Fig. 3, Table S5).

**Fig. 3.**
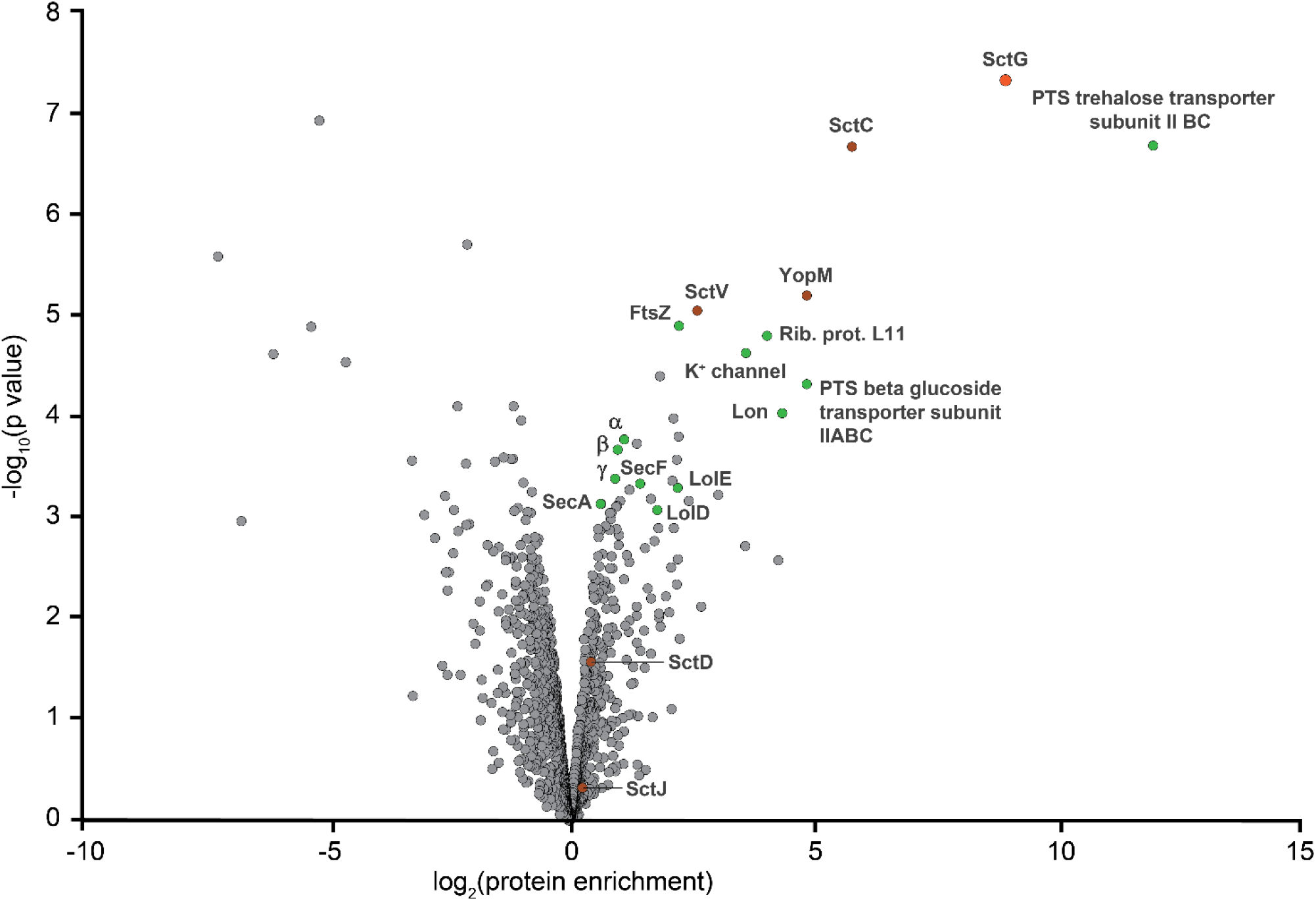
SctG interacts with T3SS components and additional non-T3SS proteins. Co-immunoprecipitation experiments with SctG-sfGFP as bait, compared to an unlabeled control strain expressing sfGFP from plasmid at comparable levels. Interactors were quantified with label-free mass spectrometry (*n*=3). The bait protein SctG is marked in orange, other T3SS components in dark red, interactors discussed in the main text in green. Greek letters denote subunits of the F1 subcomplex of the ATP synthase. A full list of all interactors with *p*<0.001 and T3SS components can be found in Table S5.

Interestingly, a number of proteins not directly linked to the T3SS were enriched highly significantly. Most prominently, the phosphoenolpyruvate (PEP)-dependent phosphotransferase system (PTS) trehalose transporter subunit EIIBC (PTS-IIBC) consistently interacted very specifically with SctG (Fig. 3), despite low overall expression levels (Table S2). A closer look at the results also showed a significant enrichment of several components of the Lol protein export pathway for the insertion of lipoproteins into the OM, namely LolD and LolE, as well as other non-T3SS proteins, such as Lon protease (termed endopeptidase La in *Y. enterocolitica*) and the prokaryotic tubulin homolog FtsZ, a major player in cell division (Fig. 3, Table S5).

Interestingly, PTS sugar transporters were shown to have a role in the regulation of the T3SS earlier [47]. We therefore constructed a clean *Δpts-IIBC* deletion mutant and tested its impact on assembly and function of the T3SS. PTS-IIBC was not required for assembly of injectisomes or effector secretion (Fig. S5a) and overexpression did not significantly impact T3SS localization or secretion itself (Fig. S5b), leaving open the functional relevance of its interaction with the pilotin. However, a proteome analysis revealed that both the pilotin SctG and OmpW were among the few proteins significantly downregulated in *Δpts-IIBC* (Table S6).

### Pilotin expression is controlled by temperature, similarly to structural T3SS components

To explain the abundant interaction of SctG with non-T3SS components, we analyzed its expression and localization in more detail. Pilotin proteins are abundant but not omnipresent in secretin-containing secretion systems [22]. Despite their consistent reported role for assembly of the highly conserved secretin, pilotins show low sequence conservation (Fig. S6a). Accordingly, we found that even the pilotin from the closely related T3SS of *P. aeruginosa* could not complement a *sctG* deletion in *Y. enterocolitica* (Fig. S6b).

Despite this low sequence conservation, pilotins have a conserved unique genetic location within the T3SS. In contrast to almost all other T3SS components, which are encoded in operons whose transcription is controlled by a common transcriptional regulator, pilotins are located in a joint operon upstream of this regulator (genetic neighborhood analysis, Fig. S7). This arrangement might lead to faster expression of the pilotin in comparison to other T3SS components, which are activated by VirF itself. In addition, the presence of an RNA thermometer between *sctG* and *virF* [42,43,48] might lead to a specific pilotin expression profile.

SctG-sfGFP was expressed at a level below the detection limit prior to the shift to 37°C, which activates the VirF-dependent expression of other T3SS components. After the temperature shift, SctG expression increased steadily over the next two hours (Fig. 4ab). Overall, the SctG-sfGFP expression profile did not differ significantly from mCherry-SctD, a basal body protein labeled in the same strain (Fig. 4b).

**Fig. 4.**
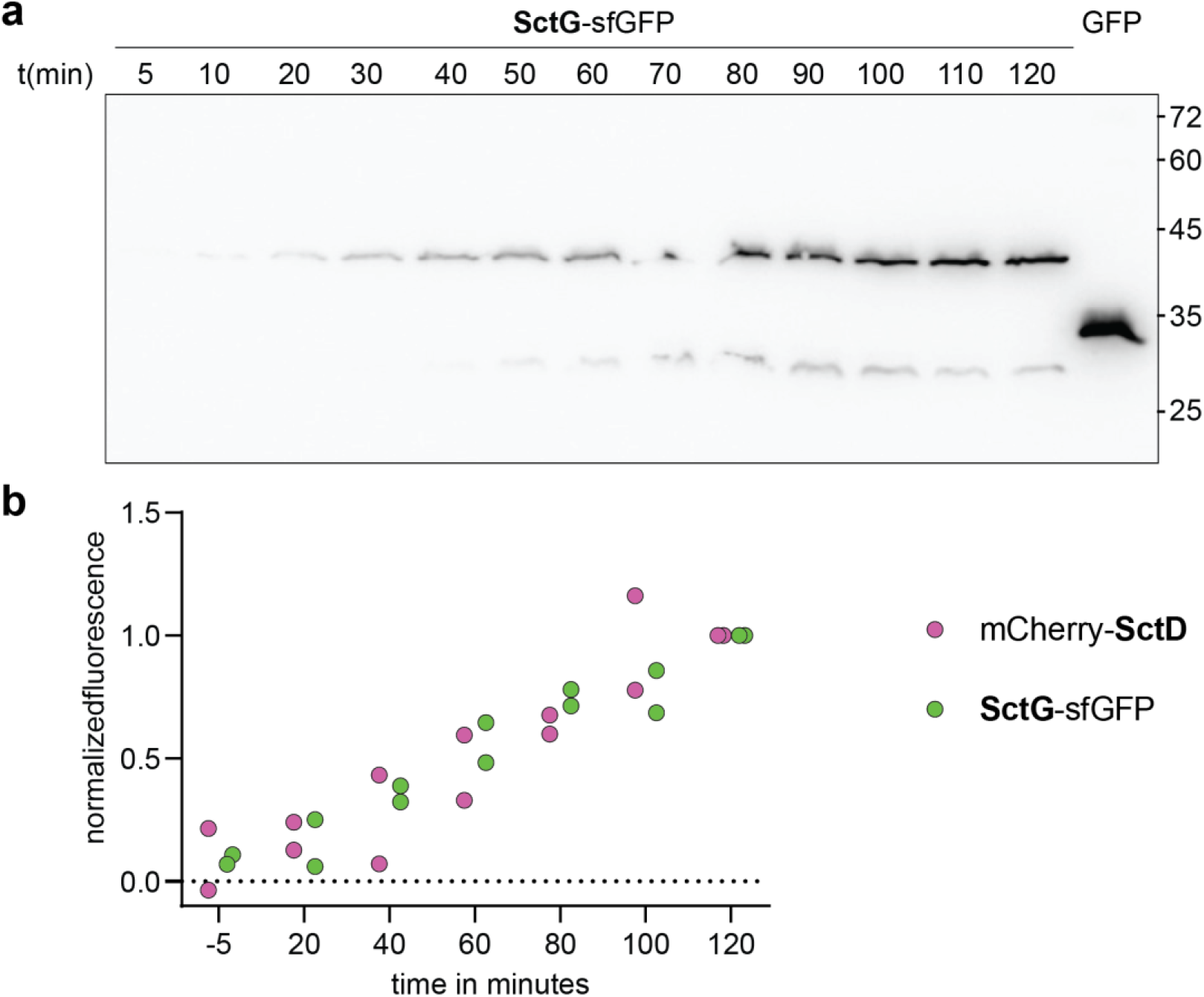
Despite its unique genetic location, SctG expression is regulated similar to other T3SS components. **a**) Expression of SctG-sfGFP (expected molecular weight 42.2 kDa) expressed from the native genetic location in *Y. enterocolitica* under non-secreting conditions. Immunoblot using antibodies directed against GFP. Control: Expression of GFP from plasmid. Numbers indicate time in min relative to temperature shift to 37°C, which triggers the expression of T3SS components. Right, molecular weight in kDa. *n*=2. **b**) Expression of SctG-sfGFP and the IM ring component mCherry-SctD, both expressed from the native genetic location in *Y. enterocolitica*, before and after temperature shift to 37°C in a double-labeled strain under non-secreting conditions. Quantification of fluorescence per bacterium, normalized by the respective 120 min value. Dots indicate individual biological replicates with a total of 52, 100, 128, 128, 217, 185, 143 bacteria (from left to right).

### Pilotins localize dynamically within the membrane and do not stably colocalize with other injectisome components

The functional SctG-sfGFP fusion allowed us to assay the cellular localization of SctG. To first test the requirement of other T3SS component or non-T3SS-related *Y. enterocolitica* proteins for SctG localization, SctG or SctG-sfGFP were expressed from plasmid in a *Y. enterocolitica* strain cured of the virulence plasmid (pYV^-^), as well as *Escherichia coli* Top10 (which has no T3SS) (Fig. S8a). While SctG-sfGFP displayed a membrane localization in *Y. enterocolitica* pYV^-^, similar to its native localization, it localized predominantly to the cytosol in *E. coli* (Fig. S8ab). Notably, expression of SctG or SctG-sfGFP negatively influenced the growth of *E. coli*, which only grew to a low OD. Microscopy experiments revealed severe lesions and plasmolysis in the respective bacteria (Fig. S8ac), indicating that the *Y. enterocolitica* pilotin mislocalizes and can thus be toxic upon ectopic expression in *E. coli*. Together, these data show that T3SS-independent species-specific bacterial factors are required for the membrane integration of the pilotin.

We then tested the localization of SctG-mCherry expressed from its native genetic location in wild-type *Y. enterocolitica* with functional T3SS. Stable stoichiometric association of the pilotin with the secretin SctC would result in the formation of membrane foci colocalizing with other injectisome components [25,44]. However, we detected a disperse membrane localization of SctG-sfGFP, which increased over time, in addition to cytosolic background fluorescence (Fig. 5a). While a certain clustering of SctG-sfGFP in the membrane was visible, these clusters did not visibly colocalize with the mCherry-labeled T3SS components SctC, SctD, or SctL, in contrast to the clear colocalization observed for mCherry-SctD and EGFP-SctQ used as positive control (Fig. 5b, Fig. S9).

**Fig. 5.**
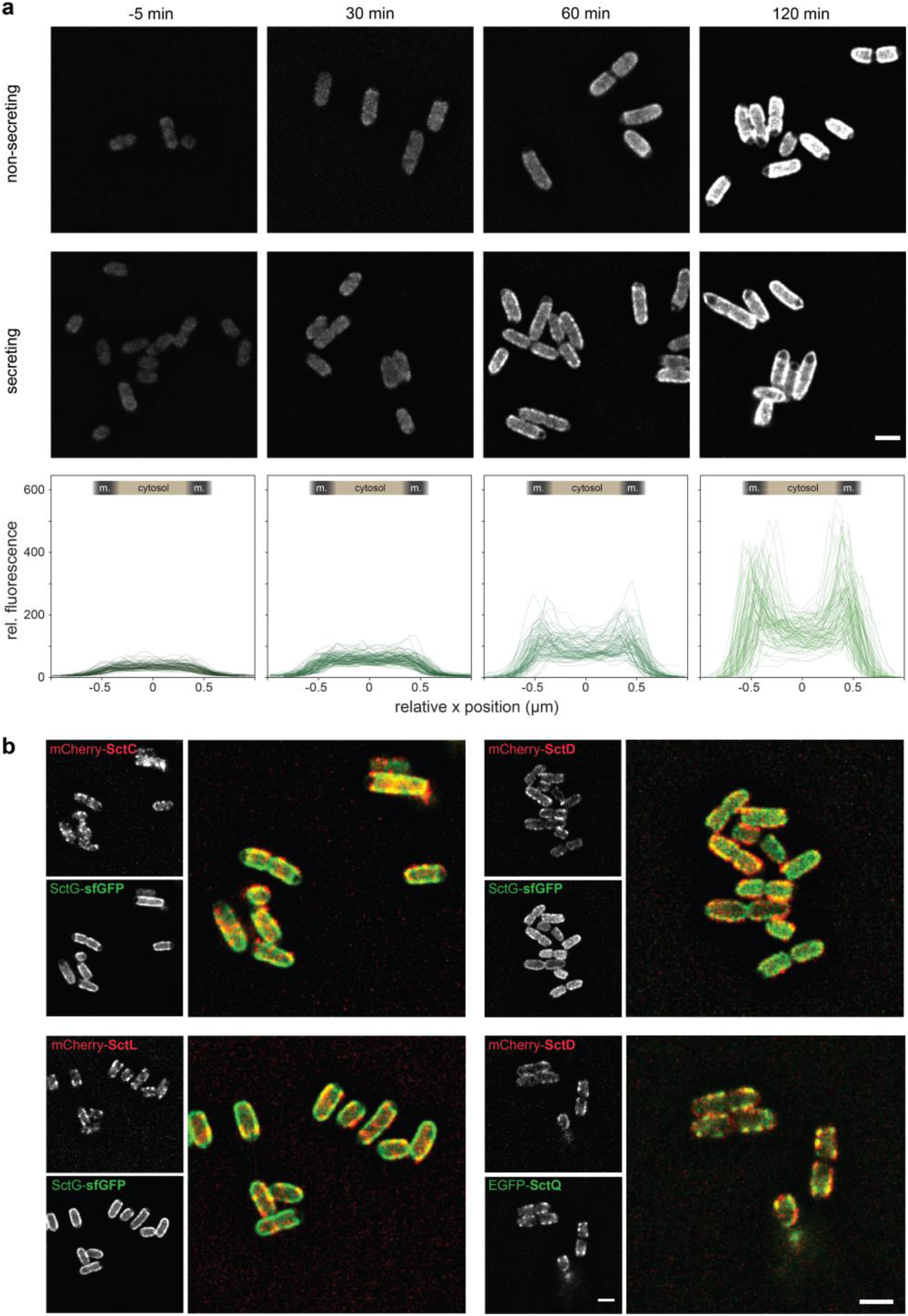
SctG predominantly localizes in the membrane, but does not significantly colocalize with other injectisome components. **a**) Time course microscopy of SctG-sfGFP in *Y. enterocolitica* under non-secreting conditions. Time with reference to the temperature shift to 37°C. Bottom, intracellular localization of SctG-sfGFP at the respective time points. Line scans across single bacteria, approximate average positions of bacterial membranes (m., dark grey) and cytosol (beige) indicated. *n*=78, 90, 81, 90 cells (left to right) from three independent biological replicates. **b**) Colocalization of labeled T3SS components (all expressed from native genetic locus on virulence plasmid) in double-labeled strains as indicated, 120 min after temperature shift to 37°C. *n*=3, except for colocalization of mCherry-SctD and EGFP-SctQ, which was reported earlier [44] and included here as a positive control (*n*=1). Scale bars = 2 μm.

To analyze the mobility and cluster formation of SctG in more detail, we tracked SctG-sfGFP in individual bacteria. Still images and overlays (Fig. 6a), as well as movies (Movie S1) showed that the localization of SctG and its clusters in the membrane strongly fluctuate, indicating diffusion within the membrane. We tested this hypothesis by fluorescence recovery after photobleaching (FRAP) experiments and found that indeed, SctG-sfGFP is highly mobile within the membrane with an average half time of recovery at the bleached pole of 21.5 s (+/-13.0s) (Fig. 6b).

**Fig. 6.**
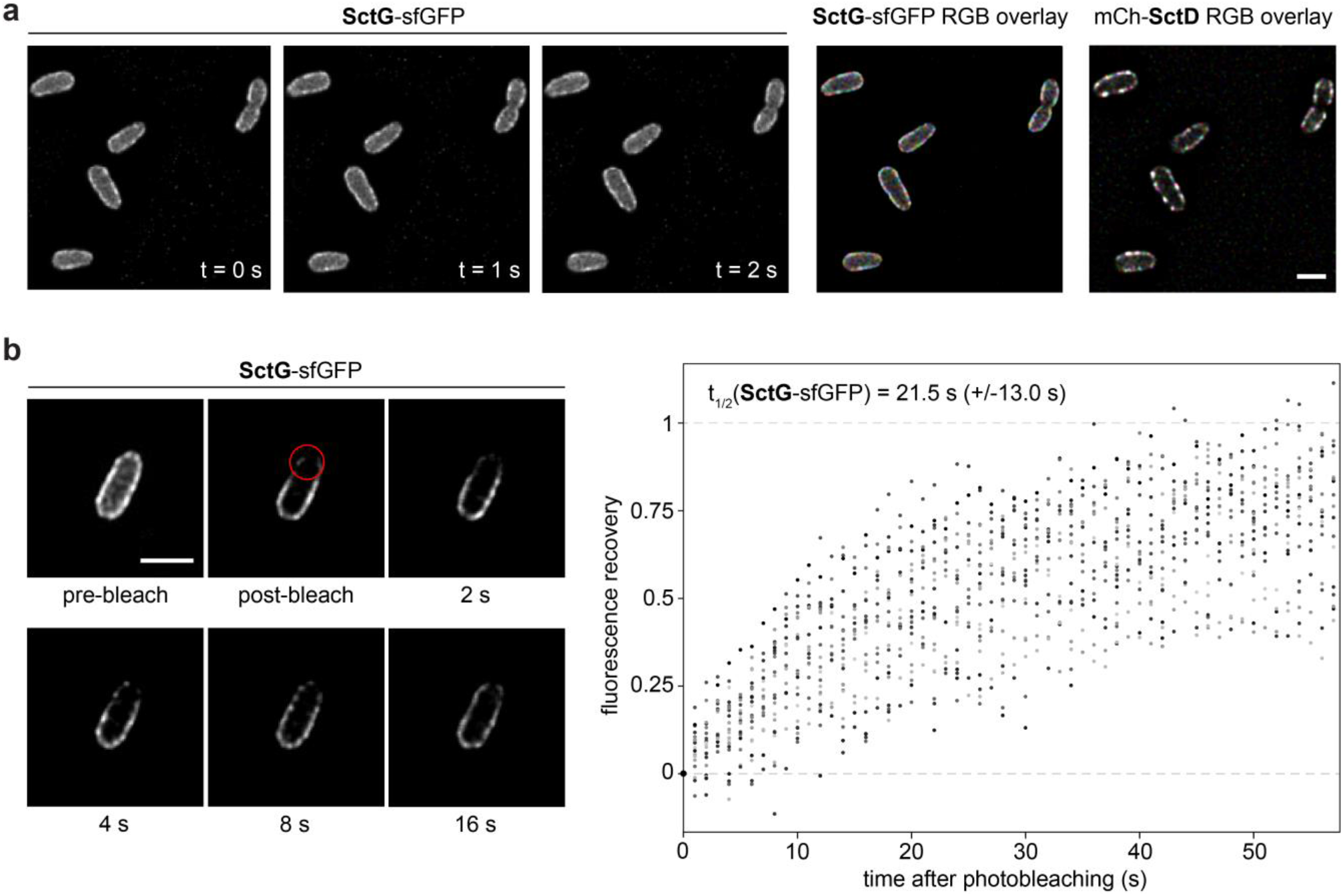
Pilotin proteins move dynamically within the membrane, forming transient patches. Fluorescence microscopy of a strain expressing SctG-sfGFP and mCherry-SctD from their native genetic locations on the virulence plasmid. **a**) Left, single micrographs of SctG-sfGFP taken with 1 s difference. Right, red/green/blue (RGB) overlay of those micrographs for SctG-sfGFP and mCherry-SctD (mCh-SctD) (*n*=3). **b**) Fluorescence recovery after photobleaching for SctG-sfGFP. Left, representative individual micrographs before and at the indicated time points after the bleach event. Bleach spot indicated by red circle. Right, quantification of fluorescence recovery after photobleaching (*n*=21 cells from 3 independent experiments, displayed in different shades of grey). The recovery half-time t1/2 was calculated for each individual recovery curve (see Material and Methods for details) and is displayed as average with standard deviation.

## Discussion

Pilotin proteins are conserved components of T3SS in many bacteria including most animal pathogenic variants. They have a unique genetic location and a central role for the assembly of the T3SS injectisome. However, we know little about their behavior and role beyond the assistance in the proper localization and oligomerization of the secretin ring of the injectisome in the OM. In this study, we confirm this role, but describe a an additional specific effect on T3SS export substrate selection. Unexpectedly, we found that pilotins are mobile membrane components, raising the possibility that they have other cellular functions after contributing to the assembly of the T3SS.

### Pilotins differentially affect major steps in T3SS assembly

Pilotins have long been known to assist the assembly of the SctC secretin ring in the OM [18,19,23,24,49–51]. Our data confirm these reports: In the absence of the pilotin, secretins localized in large foci at one of the bacterial poles (Fig. 2a). The localization of other T3SS components in absence of SctG differed from that of the secretin: SctL, a component of the cytosolic sorting platform of the T3SS, only occasionally (in 6.6% of the cells) formed single, mostly polar, foci. Similarly, we observed few (in 4.6% of the cells), also mostly polar, needles for a subset of bacteria. However, these needles and the sorting platform protein SctL did not colocalize (Fig. 2a, Fig. S2b), indicating an absence of stably assembled injectisomes. Consistent with this, the nonameric large export apparatus component SctV [52] assembled, but moved within the membrane in the absence of SctG (Fig. 2b, Fig. S2a), indicating a lack of anchoring by the ring-forming SctCDJ proteins [26]. We speculate that transient interactions of SctL and the other cytosolic T3SS components with basal body components at the poles might allow the limited export of the needle subunit. Once exported, these needles would stay stably associated with the bacteria despite the disassembly of the cytosolic sorting platform. In agreement with this hypothesis, we had found earlier that the cytosolic T3SS sorting platform components do exchange between the injectisome-bound sorting platform and a cytosolic pool [44,53] and that the sorting platform components temporarily dissociate from the remaining injectisomes under certain conditions, such as low external pH [54]. The presence of a low number of needles had previously been reported for the closely related *P. aeruginosa* T3SS [24]. Notably, the fraction of bacteria with associated needles and the overall secretion efficiency of the pilotin deletion showed an unusually high variation between individual replication experiments, suggesting an influence of external factors. However, despite our best efforts, we were unable to identify these factors.

### Role of pilotins in T3SS substrate specificity

Analysis of the T3SS secretome by SDS-PAGE and quantitative mass spectrometry revealed a specific effect of the presence of the pilotin on export of the so-called early substrates, required for formation of the needle and its tip structure (Fig. 1, Table 1). Although deletion or modifications of *sctG* have an effect on the expression of VirF and its targets (Table S2, S5), most likely due to an influence on the formation of hairpin structures in the *sctG-virF* mRNA that regulate VirF transcription as part of the thermoregulation of T3SS expression [42,43,48], the role of SctG in secretion substrate is independent from VirF levels (Fig. S1b). This leaves open the reason for the striking change of substrate specificity in absence of the pilotin, where early export substrates are much more strongly affected by the absence of SctG than the late substrates, the virulence effectors. Notably, substrate-specific effects on the secretion had been found, although not always explicitly noted, in earlier studies in *Y. enterocolitica* [15,19] and *P. aeruginosa* [24]. Taking into account the limited, probably transient interaction of the pilotin with the secretin that we observed, it is tempting to speculate that presence of the pilotin at the injectisome favors the secretion of early substrates. Subsequent dissociation of the pilotin would then contribute to a switch in export cargo specificity towards the effectors. The specific (albeit not necessarily direct) interaction of SctG with the large export apparatus component SctV (Fig. 3), which is a key player in substrate selection [55–57], supports this hypothesis. In contrast, overexpression of SctW did not increase visibly the fraction of early substrates amongst the exported proteins (Fig. S6bc).

Alternative explanations for the change in export substrate specificity might be (i) the polar localization of the secretin (and consequently any transient T3SS complex), which might change the accessibility for certain substrates, or (ii) a two-step export pathway, with a first transport step across the IM, facilitated by the T3SS export apparatus (which is assembled, but not anchored in the peptidoglycan, in the absence of the pilotin (Fig. 2b)) and a subsequent export step across the OM, facilitated by the secretin (without which no export is observed (Fig. S1)). Such a two-step transport might prevent the export of early substrates, many of which polymerize outside the cytosol. Alternatively, (iii) the at most temporary association of the cytosolic components with the membrane components (Fig. 2) could favor the export of late substrates, or (iv) strains lacking the pilotin might be able to provide less energy due to the missing link to the PTS sugar transporters or membrane stress as a consequence of secretin misassembly. If effector export is less energy costly than the export of early substrates, this might explain the observed bias in export in absence of SctG. However, in the absence of direct evidence for these alternative explanations, we consider a direct influence of SctG on the substrate specificity through interactions with other T3SS components to be the most likely explanation.

Substrate specificity switching is a central, but incompletely understood, feature type III secretion. Our data suggest that pilotin proteins play a key role in this process, together with the export apparatus components SctV and SctU [58], the gatekeeper complex [59,60] and the sorting platform [61]. If and how the interaction of the pilotin with the effector YopM (Fig. 3) contributes to this process is unclear at the moment.

### Additional interactions of the pilotin after its dissociation from the injectisome and possible functions

An unexpected finding of our study was that pilotin concentrations significantly below wild-type levels are sufficient to restore normal T3SS assembly and function (Fig. S3). Although it is one of the few T3SS machinery components encoded outside the VirF-regulated operons, native SctG expression develops very similarly to other T3SS components (Fig. 4). This is consistent with the cotranscription of *sctG* and *virF* in one operon [42] and a negligible lag phase between expression of VirF and its target proteins under the used conditions (incubation at 37° prior to activation of secretion, which is expected to fully open the *sctG-virF* RNA thermometer structure [42]). However, we found that low pilotin levels already were sufficient for injectisome assembly and secretion and that SctG does not significantly colocalize with other T3SS components (Fig. 5), but is highly mobile in the bacterial membrane (Fig. 6). This suggests that the pilotin might have additional functions beyond its role in secretin assembly, possibly after its release from the secretin. We therefore screened for additional interactors of SctG by co-immunoprecipitation.

The most striking finding of the interaction studies was the highly specific enrichment of several PTS sugar transporter subunits, most significantly the membrane-bound trehalose transporter subunit EIIBC (Fig. 3, Table S5). PTS systems are used for sugar uptake in many bacteria. They create a concentration gradient by transferring the PEP phosphoryl group onto the incoming sugar via several intermediate phosphate carriers. The EIIC subunit, which can be fused to the EIIB subunit, serves as channel in the IM. While many bacteria express several PTS transporters, PTS are often unspecific and redundant [62]. Notably, PTS were repeatedly found to act as carbohydrate sensors that control bacterial metabolism in response to the environment [63]. Transporters sensing host-specific compounds can regulate expression and activity of virulence systems [64], indicating that SctG and its main interactor, the PTS trehalose transporter subunit EIIBC, might serve to modulate T3SS function according to the bacterial environment. The fact that so far, no pilotin homologs were found in plant pathogens [3], is consistent with this hypothesis. Notably, absence of both the secretin SctG and the interacting PTS trehalose transporter subunit EIIBC led to a reduced cellular amount of OmpW, an eight-stranded beta-barrel OM protein required for resistance to phagocytosis in *E. coli* [65–67] (Table S2, S6). While the coordination of the anti-phagocytic actions of OmpW and *Y. enterocolitica* T3SS effectors appears intuitively beneficial for the bacteria during infections, OmpW itself was not found to interact with SctG and the functional interaction remains unclear for the moment.

Other significant interaction partners of the pilotin outside the T3SS are FtsZ, a prokaryotic tubulin homolog with a key role in cell division, the protease Lon, components of the ATP synthase (subunits α, β, γ), the Sec export pathway (SecF/A), and LolD/E, two components of the ABC transporter of the Lol lipoprotein export pathway in the IM. The presence of three components of the IM ATP synthase amongst the less than forty specific interactors of the pilotin protein is remarkable and, in combination with the strong interaction with the PTS sugar transporter, suggests a link with the bacterial energy metabolism. While such a link is more than conceivable, given the high energy expense of type III secretion [68], its details are unclear at the moment. As lipoproteins in the OM, pilotins are expected to interact with components of both the Sec and the Lol pathway [27,29,69]. While most models emphasize the interaction with LolA and LolB in the periplasm and OM, respectively, we detected strong binding to LolD and LolE in the IM (Fig. 3, Table S5). Of the two models proposed for the role of pilotins in the assembly of the secretin ring by Majewski and colleagues [28], our data therefore supports the second model where the pilotin is released from LolB before interacting with the secretin. An extended discussion of the interaction studies is provided in Text S1.

Taken together, our data indicate that, besides their function in the assembly of the secretin ring of the T3SS, pilotin proteins may have additional roles linked to type III secretion. In their absence, assembly of all T3SS subcomplexes and specifically the export of early secretion substrates is strongly impeded, while effectors are exported efficiently. Pilotin proteins interact with components of the T3SS, most prominently the secretin SctC and the large export apparatus component SctV, but are mobile in the bacterial membrane and do not stay attached to or even significantly colocalize with the injectisome. In line with these findings, the pilotin also interacted with other cell wall-associated proteins such as a PTS sugar transporter subunit.

A potential model integrating these findings is that during injectisome assembly, SctG interacts with SctC and facilitates its oligomerization in the OM. The presence of the pilotin supports the export of early secretion substrates, which are required for needle formation. At a later time point, SctG dissociates from the fully assembled injectisome, which then favors the export of effector proteins upon activation of the T3SS. The pilotin is mobile at this stage and may additionally support this function by interacting with other partners, including PTS sugar transporters, which could provide the necessary energy for secretion. Such species-specific functions would be in line with the observed differences between species in localization of the pilotin protein after T3SS assembly [33–37]. Challenging this model in future studies will lead to a better understanding of the role of pilotins in both assembly and function of the T3SS.

## Material and methods

### Bacterial strain generation and genetic constructs

A list of strains and plasmids used in this study can be found in Table S7. All *Y. enterocolitica* strains used in this study are based on the *Y. enterocolitica* wild-type strain MRS40 or its derivate IML421asd (ΔHOPEMTasd). In IML421asd all major virulence effector proteins (YopH,O,P,E,M,T) are absent. Furthermore, this strain harbors a deletion of the aspartate-beta-semialdehyde dehydrogenase gene, which renders the strain auxotrophic for diaminopimelic acid (DAP), making it suitable for work in an biosafety class 1 environment [70].

### Bacterial cultivation, secretion assays and protein analysis

*Y. enterocolitica* day cultures were inoculated from fresh stationary overnight cultures supplemented with nalidixic acid (35 mg/ml) and DAP (80 mg/ml), where required, to an OD_600_ of 0.15 for secreting and 0.12 for non-secreting conditions, respectively. To select for the maintenance of expression plasmids, ampicillin was added (0.2 mg/ml). As rich growth medium, BHI (brain heart infusion broth) supplemented with DAP (80 mg/ml) where required, glycerol (0.4%) and MgCl2 (20 mM) was used. For secreting conditions, 5 mM EGTA were added; for non-secreting conditions, 5 mM CaCl2 were added to the pre-warmed (approx. 55°C) medium, which was filtered through a 0.45 μm filter before the addition of other supplements and cooled down to ambient temperature for the experiments. After inoculation, day cultures were incubated at 28°C for 90 min to reach exponential growth phase. Expression of the *yop* regulon was then induced by a rapid temperature shift to 37°C in a water bath. Where indicated, protein expression from plasmid (derivates of pBAD-His/B (Invitrogen) in all cases except expression of VirF from pCL5 [71]) was induced at this point by the addition of 0.02–1.0% L-arabinose as indicated for the respective experiments.

For protein secretion assays and analysis of total cellular proteins, bacteria were incubated for 2-3 h at 37°C, as indicated. 2 ml of the culture were collected at 21,000 *g* for 10 min. The supernatant was removed from the total cell pellet and proteins were precipitated by addition of a final concentration of 10% trichloroacetic acid (TCA) and incubation at 4°C for 1–8 h. Precipitated proteins were collected by centrifugation for 15 min at 21,000 *g* and 4°C, the pellet was washed once with 1 ml ice cold acetone and subsequently resuspended and normalized in SDS-PAGE loading buffer. Total cellular protein samples were normalized to 0.3 OD units (ODu, 1 ODu is equivalent to 1 ml of culture at on OD_600_ of 1, corresponding to approximately 5 x 10^8^ *Y. enterocolitica*) and supernatant samples to the equivalent 0.6 ODu. Before loading, samples were incubated for 5 min at 99°C. Separation was performed on 11%, 15% or gradient 12–20% SDS-PAGE gels, using BlueClassic Prestained Marker (Jena Biosciences) as a size standard. For visualization, the SDS-PAGE gels were stained with InstantBlue (Expedeon). For immunoblots, the separated proteins were transferred from the SDS-PAGE gel onto a nitrocellulose membrane. Primary mouse antibodies against GFP (ThermoFisher Proteintech 66002-1-lg, 1:4000) were used in combination with secondary anti-mouse antibodies conjugated to horseradish peroxidase (GE Healthcare NXA931, 1:5000). For visualization, ECL chemiluminescence substrate (Pierce) was used in a LAS-4000 Luminescence Image Analyzer.

### Fluorescence microscopy

For fluorescence microscopy, bacteria were treated as described above. After 2–3 h at 37°C, 400 μl of bacterial culture were collected by centrifugation (2400 × g, 2 min) and reconstituted in 200 μl minimal medium (100 mM 2-[4-(2-Hydroxyethyl)piperazin-1-yl]ethane-1-sulfonic acid (HEPES) pH 7.2, 5 mM (NH_4_)_2_SO_4_, 100 mM NaCl, 20 mM sodium glutamate, 10 mM MgCl_2_, 5 mM K_2_SO_4_, 50 mM 2-(N-morpholino)ethanesulfonic acid (MES), 50 mM glycine). 2 μl of resuspension were spotted on agarose pads (1.5% low melting agarose (Sigma-Aldrich) in minimal medium) in glass depression slides (Marienfeld). Where required, agarose pads and media were supplemented with 80 mg/ml DAP for ΔHOPEMTasd-based strains, 5 mM CaCl_2_ for non-secreting conditions, or 5 mM EGTA for secreting conditions.

Microscopy was performed on a Deltavision Elite Optical Sectioning Microscope equipped with an UPlanSApo 100×/1.40 oil objective (Olympus), using an Evolve EMCCD camera (Photometrics). Where applicable, the prepared samples were illuminated first in the mCherry = 0.4 s with a mCherry filter set and afterwards in GFP = 0.2 sec with a GFP filter set. Depending on the experiment, z stacks with 1-15 slices (Δz = 0.15 μm) per fluorescence channel were acquired. The acquired micrographs where subsequently deconvolved using softWoRx 7.0.0 (standard “conservative” settings). Images were further processed with FIJI (ImageJ 1.51f/1.52i/1.52n) [72]. Where necessary, a drift correction with the StackReg Plugin [73] was performed. Fluorescence quantification was performed in FIJI. For presentation of micrographs in a figure, representative fields of view were selected, and brightness and contrast of the micrographs was adjusted identically within the compared image sets.

### Maleimide-based needle (SctF) staining

Maleimide-based staining to visualize the injectisome needles was performed as described in [54]. Briefly, expression of SctF_S5C_ from the pBAD-His/B plasmid was induced by addition of 1.0% L-arabinose at the time of the temperature shift in strains expressing the native SctF at the same time. After 2-3 h, bacteria were collected and resuspended in 0.2 volumes of minimal medium supplemented with 5 μM CF 488A maleimide dye (Sigma-Aldrich, USA) at 37 °C for 5 min. After staining, cells were washed once with 1500 μl of minimal medium and resuspended in 200 μl of minimal medium. 2 μl of bacterial suspension were spotted on 1.5% agarose pads in minimal medium and visualized under the microscope as described above.

### Classification and quantification of fluorescent foci

To determine the number of stable SctV-EGFP foci, seven micrographs were taken in 15 s intervals and the minimal intensity for each location was determined from the deconvoluted micrographs. Foci were then identified in FIJI using an intensity threshold of 700 (determined in images of the wild-type background strain) and the “analyze particles” function with a minimal size of 4 pixels and a particle circularity of 0.5-1.0. To evaluate the fraction of bacteria with foci for SctC-mCherry, mCherry-SctL and SctF_S5C_ in the absence of SctG, foci were manually identified and classified as polar, subpolar and lateral based on their subcellular localization in deconvoluted micrographs. In case a cell displayed multiple foci in different localization classes, bacteria were classified as containing a lateral, subpolar, polar focus in descending order (e.g. a bacterium with a lateral and a polar focus was assigned “lateral”). The overall number of bacteria in a field of view was determined by using the deep learning based network, DeepBacs [74], that was previously trained with differential interferometry contrast (DIC) micrographs of *Y. enterocolitica*. The overall percentage of cells with foci is displayed in the figure.

### Fluorescence distribution analysis

For analysis of the fluorescence distribution across bacteria, line scan analysis was performed on deconvoluted micrographs in FIJI. With the straight line freehand tool, measurements areas were defined from one lateral membrane to the other at a 90° angle. Per condition, 3 fields of view with 18-30 cells were analyzed and the background was measured. The line scans were individually corrected for background fluorescence, normalized (fluorescence fraction relative to overall fluorescence over background) and centered at the position closest to center of the total fluorescence over background.

### Fluorescence recovery after photobleaching (FRAP)

Cells were grown in non-secreting medium as described above. Samples were taken after 3 h at 37°C. To minimize unnecessary bleaching, no z stacks were acquired. Three pre-bleach images were acquired and locations near selected bacterial cell poles were photobleached by individual 30 ns pulses of a 488 nm laser. Recovery of the bleached foci was followed by time-lapse microscopy. Deconvolved micrographs were used for the fluorescence quantification.

The relative fluorescence of the bleached spot was calculated as follows:

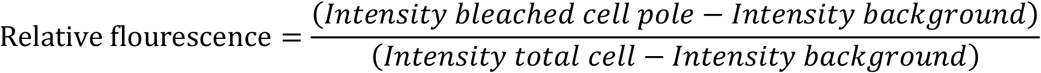

The relative fluorescence was then normalized (average pre-bleach value set to 1; post-bleach value set to 0) and an exponential recovery was fitted using the equation:

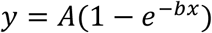

where y stands for the relative fluorescence, x for the time, A for the overall ratio of fluorescence recovery, and b for the inverse of the time constant τ_1/2_. For the fit, A was restricted to values between 0.7 and 1.1. Each bleach curve was fitted individually, and fits with R^2^ < 0.7 were excluded from further analysis. The half-time of recovery was calculated from the time constant as follows:

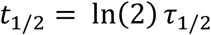

### Proteome and secretome analysis by shotgun proteomics based mass spectrometry

Strains were grown and prepared as described above. After 2.5 h at 37°, cultures were normalized to an OD_600_ of 0.5, and 2 ml were harvested by centrifugation at 9,391 *g* for 2 min. Cell were washed 3 times with 1x phosphate buffered saline (PBS) (8 g/l NaCl, 0.2 g/l KCl, 1.78 g/l Na_2_HPO_4_*2H_2_O, 0.24 g/l KH_2_PO_4_, pH 7.4). After washing, the cell pellet was resuspended in 300 μl lysis buffer (2% sodium lauroyl sarcosinate (SLS), 100 mM ammonium bicarbonate (ABC)) and incubated at 90°C for 10 min. Protein concentrations were measured using a BCA assay (Thermo Scientific) and following reduction in 5 mM Tris(2-carboxyethyl)phosphin (TCEP, 15 min at 90 °C) and alkylation in 10 mM iodoacetamid (30 min, 20°C in the dark), 50 μg protein was used for tryptic digestion (1 μg, Serva) at 30°C overnight in presence of 0.5% SLS. After digestion, SLS was precipitated by adding a final concentration of 1.5% trifluoroacetic acid (TFA).

Desalting of cleared peptide samples was performed in C18 solid phase extraction (SPE) cartridges (Macherey-Nagel). First, cartridges were prepared by adding acetonitrile, followed by equilibration using 0.1% TFA and loading of peptide samples. Cartridges were washed with buffer containing 5% acetonitrile and 0.1% TFA, and finally eluted with 50% acetonitrile and 0.1% TFA into a fresh tube. Eluted peptides were dried and reconstituted in 0.1% TFA for LC-MS analysis.

Purified peptides were analyzed by liquid chromatography-mass spectrometry (MS) carried out on a Q-Exactive Plus instrument connected to an Ultimate 3000 rapid-separation liquid chromatography (RSLC) nano instrument and a nanospray flex ion source (all Thermo Scientific). Peptide separation was performed on a reverse-phase high-performance liquid chromatography (HPLC) column (75 μm by 42 cm) packed in-house with C_18_ resin (2.4 μm; Dr. Maisch GmbH). The peptides were first loaded onto a C18 precolumn (preconcentration set-up) and then eluted in backflush mode with a gradient from 98 % solvent A (0.15 % formic acid) and 2 % solvent B (99.85 % acetonitrile, 0.15 % formic acid) to 25 % solvent B over 66 min, continued from 25 to 40 % of solvent B up to 90 min. The flow rate was set to 300 nl/min. The data acquisition mode for label-free quantification (LFQ) was set to obtain one high-resolution MS scan at a resolution of 70,000 (*m/z* 200) with scanning range from 375 to 1500 *m/z* followed by MS/MS scans of the 10 most intense ions at a resolution of 17,500. To increase the efficiency of MS/MS shots, the charged state screening modus was adjusted to exclude unassigned and singly charged ions. The dynamic exclusion duration was set to 10 s. The ion accumulation time was set to 50 ms (both MS and MS/MS). The automatic gain control (AGC) was set to 3 × 10^6^ for MS survey scans and 1 × 10^5^ for MS/MS scans.

Due to an instrument upgrade, the QExactive Plus system was replaced by an Exploris 480 mass spectrometer (Thermo Scientific). For total proteome analysis, purified peptides were analyzed using identical LC settings, and the Exploris MS data acquisition settings were the following: one high-resolution MS scan at a resolution of 60,000 (m/z 200) was obtained with scanning range from 350 to 1650 m/z followed by MS/MS scans of 2 sec cycle time at a resolution of 15,000. Charge state inclusion was set between 2 and 6. The dynamic exclusion duration was set to 14 s. The ion accumulation time was set to 25 ms for MS and AUTO for MS/MS. The automatic gain control (AGC) was set to 300% for MS survey scans and 200% for MS/MS scans.

LFQ analysis was performed using MaxQuant [75] in standard settings using a *Y. enterocolitica* protein database containing proteins of the closely related *Y. enterocolitica* strain W22703 [76] and of the pYVe227 virulence plasmid (GenBank entry AF102990.1). Statistical follow-up analysis of the MaxQuant LFQ data was performed on an updated SafeQuant [77,78] R-script modified to routinely process MaxQuant “protein groups” outputs. The missing value imputation was adapted from Perseus [79] in default settings and implemented into the update SafeQuant script. Unless otherwise indicated, proteins with ≥3 detected peptides and a *p* value ≤ 0.01 of the sample versus the control using Student’s t-test were considered as enriched and displayed in the tables. Intensity values were shaded according to their value (blue for high intensity; transparent for low intensity) for easier visualization.

For secretome analysis and quantification of secreted effectors in minimal medium, 5 ml BHI cultures were grown under secreting conditions in 15 ml tubes as otherwise described before. After two hours of incubation at 37°C, the medium was changed from BHI to minimal medium supplemented with nalidixic acid (35 mg/ml), diaminopimelic acid (80 mg/ml) where required, glycerol (0.4%), MgCl_2_ (20 mM) and EGTA (5 mM). Next, bacteria were again incubated at 37°C shaking for one hour. Samples were OD-normalized in a 2 ml volume, and bacteria were removed by centrifugation (10 min at 21,000 *g*, 4°C). In the next step, the supernatant was removed and precipitated with a final concentration of 10% TCA for 1–8 h at 4°C; proteins were collected by centrifugation for 15 min at 21,000 *g* at 4°C and washed once with 1 ml ice cold acetone. A detailed description of the experimental procedure is shown in [80]. The remaining pellet was resuspended in 300 μl lysis buffer and incubated at 90°C for 10 min. Following reduction and alkylation (see above), proteins were digested using 1 μg trypsin. All further steps including C18-SPE, LC-MS analysis and LFQ data analysis were identical to the ones described above. The LC separating gradient length was reduced to 45 min due to lower sample complexity.

### Co-immunoprecipitation and shotgun proteomics

For co-immunoprecipitation, 100 ml of non-secreting culture medium were inoculated to an OD_600_ of ~0.15 and treated as described above. After 2.5 h incubation at 37°C, the cultures were transferred to 50 ml tubes and centrifuged at 2,000 *g* for 15 min at 4°C. Pellets were washed with 10 ml of 1x PBS. The contents of two tubes were pooled and centrifuged at 2,000 *g* for 15 min at 4°C. The pellet was resuspended in 2 ml HNN lysis buffer (50 mM HEPES pH 7.5, 150 mM NaCl, 50mM NaF, sterile filtered; protease inhibitor (cOmplete Mini, EDTA free (Roche)) added before use), and incubated for 30 min on a turning wheel at 4°C. Afterwards the bacteria were sonicated with 12 pulses à 30 s in a Hielscher UP200ST ultrasonic homogenizer. The cells were again placed on a turning wheel for 30 min at 4°C and the sonification process and the subsequent incubation on a turning wheel were repeated. Unbroken cells were removed by centrifugation (20,000 *g*, 20 min, 4°C). For affinity purification, 10 μl of bead slurry (GFP-Trap Magnetic Agarose, Chromotek) were added to the lysate and incubated shaking for 1 h at 4°C on a turning wheel. Beads were washed with 500 μl HNN lysis buffer + 0.1% NP40, followed by seven consecutive washing steps with 700 μl 100 mM ABC. 200 μl of elution buffer (100 mM ABC, 1 μg trypsin per sample) were added to the beads and incubated for 45 min on a thermomixer at 30°C at 1200 rpm. Beads were separated and the supernatant was collected. Beads were washed twice using 80 μl of elution buffer 2 (1 M urea, 100 mM ABC, 5 mM TCEP), combined with the first eluate fraction and incubated o/n at 30°C to complete proteolysis. To monitor the experimental steps, samples for Western blot analysis were taken throughout the whole procedure. After digest, 10 mM iodoacetamid were added and the samples were incubated for 30 min in the dark. For desalting, C18 solid phase extraction cartridges samples were acidified and the procedure was carried out as described above. The samples were analyzed on a QExactive Plus mass spectrometer using a resolving LC gradient of 40 min. The analytical settings and data analysis steps including MaxQuant and SafeQuant are described above. Unless otherwise indicated, proteins with ≥3 detected peptides and a *p* value ≤ 0.001 of the sample versus the control using Student’s t-test were considered as enriched and displayed in the tables.

### Sequence analysis

For the analysis of the genetic environment of T3SS pilotin proteins from model organisms, the following GenBank sequences were used: AF102990.1 (*Yersinia enterocolitica* plasmid pYVe227, complete sequence), NC_002516.2 (*Pseudomonas aeruginosa* PAO1, complete genome reference sequence), NC_003197.2 (*Salmonella enterica* subsp. *enterica* serovar Typhimurium str. LT2, complete genome reference sequence), NC_004851.1 (*Shigella flexneri* 2a str. 301 plasmid pCP301, complete sequence), AM286415.1 (*Yersinia enterocolitica* subsp. *enterocolitica* 8081, complete genome). Multiple sequence alignment of the *sctG* and *sctC* genes of the same systems was performed using T-COFFEE 11.0 [81].

## Acknowledgements

This work was supported by the Max Planck Society. The authors thank Christoph Spahn (Max Planck Institute for Terrestrial Microbiology, Marburg, Germany) for fruitful discussions about microscopy data analysis and his help with implementing the DeepBacs segmentation tool for *Y. enterocolitica*.

## Author contributions

S. W. performed the majority of the experiments and data analysis and participated in study design and writing of the manuscript. M.F. performed experiments, implemented methods and generated key strains for this study. C.H., C.B. and K.L. assisted in experiments and strain generation. W.S., G.A. T. G. from the Proteomics facility provided this technology and assisted in data evaluation and statistical analysis. A.D. conceived and designed the study, performed data analysis and wrote the manuscript.

## Footnotes

1 The T3SS is at the core of both injectisomes and bacterial flagella [11,82]. In this manuscript, “T3SS” refers to the injectisome-type, virulence-associated system.

2 This manuscript uses the common nomenclature for T3SS components [83,84]; the species-specific nomenclature of the *Yersinia* pilotin is YscW; other names can be found in Table S1.

## References

1. Galán JE, Wolf-Watz H. Protein delivery into eukaryotic cells by type III secretion machines. Nature. 2006;444(7119):567–73.

2. Cornelis GR. The type III secretion injectisome. Nat Rev Microbiol. 2006;4(11):811–25.

3. Büttner D. Protein export according to schedule: architecture, assembly, and regulation of type III secretion systems from plant- and animal-pathogenic bacteria. Microbiol Mol Biol Rev. 2012;76(2):262–310.

4. Burkinshaw BJ, Strynadka NC. Assembly and structure of the T3SS. Biochim Biophys Acta - Mol Cell Res. 2014;1843(8):1648–93.

5. Wagner S, Grin I, Malmsheimer S, Singh N, Torres-Vargas CE, Westerhausen S. Bacterial type III secretion systems: a complex device for the delivery of bacterial effector proteins into eukaryotic host cells. FEMS Microbiol Lett. 2018;365(19).

6. Diepold A, Wagner S. Assembly of the bacterial type III secretion machinery. FEMS Microbiol Rev. 2014;38(4):802–22.

7. Journet L, Agrain C, Broz P, Cornelis GR. The needle length of bacterial injectisomes is determined by a molecular ruler. Science (80-). 2003;302(5651):1757–60.

8. Wee DH, Hughes KT. Molecular ruler determines needle length for the Salmonella Spi-1 injectisome. Proc Natl Acad Sci. 2015;112(13):4098–103.

9. Chakravortty D, Rohde M, Jäger L, Deiwick J, Hensel M. Formation of a novel surface structure encoded by Salmonella Pathogenicity Island 2. EMBO J. 2005;24(11):2043–52.

10. Blocker AJ, Gounon P, Larquet E, Niebuhr K, Cabiaux V, Parsot C, et al. The tripartite type III secreton of Shigella flexneri inserts IpaB and IpaC into host membranes. J Cell Biol. 1999;147(3):683–93.

11. Diepold A, Armitage JP. Type III secretion systems: the bacterial flagellum and the injectisome. Philos Trans R Soc B Biol Sci. 2015;370(1679):20150020.

12. Kudryashev M, Diepold A, Amstutz M, Armitage JP, Stahlberg H, Cornelis GR. Yersinia enterocolitica type III secretion injectisomes form regularly spaced clusters, which incorporate new machines upon activation. Mol Microbiol. 2015;95(5):875–84.

13. Zhang Y, Lara-Tejero M, Bewersdorf J, Galán JE. Visualization and characterization of individual type III protein secretion machines in live bacteria. Proc Natl Acad Sci U S A. 2017;114(23):6098–103.

14. Burgess JL, Case HB, Burgess RA, Dickenson NE. Dominant negative effects by inactive Spa47 mutants inhibit T3SS function and Shigella virulence. PLoS One. 2020;15(1):e0228227.

15. Allaoui A, Scheen R, Lambert de Rouvroit C, Cornelis GR. VirG, a Yersinia enterocolitica lipoprotein involved in Ca2+ dependency, is related to exsB of Pseudomonas aeruginosa. J Bacteriol. 1995;177(15):4230–7.

16. Hardie KR, Lory S, Pugsley AP. Insertion of an outer membrane protein in Escherichia coli requires a chaperone-like protein. EMBO J. 1996;15(5):978–88.

17. Nouwen N, Ranson N, Saibil HR, Wolpensinger B, Engel A, Ghazi A, et al. Secretin PulD: association with pilot PulS, structure, and ion-conducting channel formation. Proc Natl Acad Sci U S A. 1999;96(14):8173–7.

18. Schuch R, Maurelli AT. MxiM and MxiJ, base elementsn of the Mxi-Spa type III secretion system of Shigella, interact with and stabilize the MxiD secretin in the cell envelope. J Bacteriol. 2001;183(24):6991–8.

19. Burghout P, Beckers F, de Wit E, van Boxtel R, Cornelis GR, Tommassen J, et al. Role of the pilot protein YscW in the biogenesis of the YscC secretin in Yersinia enterocolitica. J Bacteriol. 2004;186(16):5366–75.

20. Korotkov K V, Gonen T, Hol WGJ. Secretins: dynamic channels for protein transport across membranes. Trends Biochem Sci. 2011;

21. Gu S, Rehman S, Wang X, Shevchik VE, Pickersgill RW. Structural and functional insights into the pilotin-secretin complex of the type II secretion system. PLoS Pathog. 2012;8(2).

22. Koo J, Burrows LL, Lynne Howell P. Decoding the roles of pilotins and accessory proteins in secretin escort services. FEMS Microbiol Lett. 2012;328(1):1–12.

23. Rau R, Darwin AJ. Identification of YsaP, the pilotin of the Yersinia enterocolitica Ysa type III secretion system. J Bacteriol. 2015;197(17):2770–9.

24. Perdu C, Huber P, Bouillot S, Blocker AJ, Elsen S, Attree I, et al. ExsB is required for correct assembly of Pseudomonas aeruginosa type III secretion apparatus in the bacterial membrane and full virulence in vivo. Infect Immun. 2015;IAI.00048-15.

25. Diepold A, Amstutz M, Abel S, Sorg I, Jenal U, Cornelis GR. Deciphering the assembly of the Yersinia type III secretion injectisome. EMBO J. 2010;29(11):1928–40.

26. Diepold A, Wiesand U, Cornelis GR. The assembly of the export apparatus (YscR,S,T,U,V) of the Yersinia type III secretion apparatus occurs independently of other structural components and involves the formation of an YscV oligomer. Mol Microbiol. 2011;82(2):502–14.

27. Okon M, Moraes TF, Lario PI, Creagh AL, Haynes CA, Strynadka NC, et al. Structural Characterization of the Type-III Pilot-Secretin Complex from Shigella flexneri. Structure. 2008;16(10):1544–54.

28. Majewski DD, Okon M, Heinkel F, Robb CS, Vuckovic M, McIntosh LP, et al. Characterization of the Pilotin-Secretin Complex from the Salmonella enterica Type III Secretion System Using Hybrid Structural Methods. Structure. 2021;29(2):125–138.e5.

29. Collin S, Guilvout I, Nickerson NN, Pugsley AP. Sorting of an integral outer membrane protein via the lipoprotein-specific Lol pathway and a dedicated lipoprotein pilotin. Mol Microbiol. 2011;80(3):655–65.

30. Chami M, Guilvout I, Gregorini M, Rémigy HW, Müller SA, Valerio M, et al. Structural insights into the secretin PulD and its trypsin-resistant core. J Biol Chem. 2005;280(45):37732–41.

31. Yin M, Yan Z, Li X. Structural insight into the assembly of the type II secretion system pilotin–secretin complex from enterotoxigenic Escherichia coli. Nat Microbiol. 2018;3(5):581–7.

32. Chernyatina AA, Low HH. Core architecture of a bacterial type II secretion system. Nat Commun. 2019;10(1):1–10.

33. Hu B, Morado DR, Margolin W, Rohde JR, Arizmendi O, Picking WL, et al. Visualization of the type III secretion sorting platform of Shigella flexneri. Proc Natl Acad Sci. 2015;112(4):1047–52.

34. Flacht L, Lunelli M, Kaszuba K, Chen ZA, O’ Reilly FJ, Rappsilber J, et al. Integrative structural analysis of the type III secretion system needle complex from Shigella flexneri. Protein Sci. 2023;(September 2022):1–15.

35. Hu B, Lara-Tejero M, Kong Q, Galán JE, Liu J. In Situ Molecular Architecture of the Salmonella Type III Secretion Machine. Cell. 2017;168(6):1065–1074.e10.

36. Hu J, Worrall LJ, Hong C, Vuckovic M, Atkinson CE, Caveney N, et al. Cryo-EM analysis of the T3S injectisome reveals the structure of the needle and open secretin. Nat Commun. 2018;9(1):3840.

37. Miletic S, Fahrenkamp D, Goessweiner-Mohr N, Wald J, Pantel M, Vesper O, et al. Substrate-engaged type III secretion system structures reveal gating mechanism for unfolded protein translocation. Nat Commun. 2021;12(1):1546.

38. Troisfontaines P, Cornelis GR. Type III Secretion: More Systems Than You Think. Physiology. 2005;20(5):326–39.

39. Abby SS, Rocha EPC. The Non-Flagellar Type III Secretion System Evolved from the Bacterial Flagellum and Diversified into Host-Cell Adapted Systems. PLoS Genet. 2012;8(9):e1002983.

40. Iriarte M, Cornelis GR. Identification of SycN, YscX, and YscY, three new elements of the Yersinia yop virulon. J Bacteriol. 1999;181(2):675–80.

41. Diepold A, Wiesand U, Amstutz M, Cornelis GR. Assembly of the Yersinia injectisome: the missing pieces. Mol Microbiol. 2012;85(5):878–92.

42. Böhme K, Steinmann R, Kortmann J, Seekircher S, Heroven AK, Berger E, et al. Concerted actions of a thermo-labile regulator and a unique intergenic RNA thermosensor control Yersinia virulence. PLoS Pathog. 2012;8(2):e1002518.

43. Schwiesow L, Lam H, Dersch P, Auerbuch V. Yersinia Type III Secretion System Master Regulator LcrF. J Bacteriol. 2016;198(4):604–14.

44. Diepold A, Sezgin E, Huseyin M, Mortimer T, Eggeling C, Armitage JP. A dynamic and adaptive network of cytosolic interactions governs protein export by the T3SS injectisome. Nat Commun. 2017;8(1):15940.

45. Milne-Davies B, Helbig C, Wimmi S, Cheng DWC, Paczia N, Diepold A. Life After Secretion—Yersinia enterocolitica Rapidly Toggles Effector Secretion and Can Resume Cell Division in Response to Changing External Conditions. Front Microbiol. 2019;10:2128.

46. Kaniga K, Delor I, Cornelis GR. A wide-host-range suicide vector for improving reverse genetics in Gram-negative bacteria: inactivation of the blaA gene of Yersinia enterocolitica. Gene. 1991;109(1):137–41.

47. Mazé A, Glatter T, Bumann D. The central metabolism regulator EIIAGlc switches salmonella from growth arrest to acute virulence through activation of virulence factor secretion. Cell Rep. 2014;7(5):1426–33.

48. Hoe N, Goguen JD. Temperature sensing in Yersinia pestis: translation of the LcrF activator protein is thermally regulated. J Bacteriol. 1993;175(24):7901–9.

49. Koster M, Bitter W, de Cock H, Allaoui A, Cornelis GR, Tommassen J. The outer membrane component, YscC, of the Yop secretion machinery of Yersinia enterocolitica forms a ring-shaped multimeric complex. Mol Microbiol. 1997;26(4):789–97.

50. Daefler S, Russel M. The Salmonella typhimurium InvH protein is an outer membrane lipoprotein required for the proper localization of InvG. Mol Microbiol. 1998;28(6):1367–80.

51. Crago A, Koronakis V. Salmonella InvG forms a ring-like multimer that requires the InvH lipoprotein for outer membrane localization. Mol Microbiol. 1998;30(1):47–56.

52. Abrusci P, Vergara-Irigaray M, Johnson S, Beeby MD, Hendrixson DR, Roversi P, et al. Architecture of the major component of the type III secretion system export apparatus. Nat Struct Mol Biol. 2013;20(1):99–104.

53. Diepold A, Kudryashev M, Delalez NJ, Berry RM, Armitage JP. Composition, Formation, and Regulation of the Cytosolic C-ring, a Dynamic Component of the Type III Secretion Injectisome. PLOS Biol. 2015;13(1):e1002039.

54. Wimmi S, Balinovic A, Jeckel H, Selinger L, Lampaki D, Eisemann E, et al. Dynamic relocalization of cytosolic type III secretion system components prevents premature protein secretion at low external pH. Nat Commun. 2021;12(1):1625.

55. Bange G, Kümmerer N, Engel C, Bozkurt G, Wild K, Sinning I. FlhA provides the adaptor for coordinated delivery of late flagella building blocks to the type III secretion system. Proc Natl Acad Sci U S A. 2010;107(25):11295–300.

56. Xing Q, Shi K, Portaliou AG, Rossi P, Economou A, Kalodimos CG. Structures of chaperone-substrate complexes docked onto the export gate in a type III secretion system. Nat Commun. 2018;9(1):1773.

57. Inoue Y, Kinoshita M, Kida M, Takekawa N, Namba K, Imada K, et al. The FlhA linker mediates flagellar protein export switching during flagellar assembly. Commun Biol. 2021;4(1).

58. Sorg I, Wagner S, Amstutz M, Müller SA, Broz P, Lussi Y, et al. YscU recognizes translocators as export substrates of the Yersinia injectisome. EMBO J. 2007;26(12):3015–24.

59. Day JB, Plano G V. A complex composed of SycN and YscB functions as a specific chaperone for YopN in Yersinia pestis. Mol Microbiol. 1998;30(4):777–88.

60. Botteaux A, Sory M-P, Biskri L, Parsot C, Allaoui A. MxiC is secreted by and controls the substrate specificity of the Shigella flexneri type III secretion apparatus. Mol Microbiol. 2009;71(2):449–60.

61. Lara-Tejero M, Kato J, Wagner S, Liu X, Galán JE. A Sorting Platform Determines the Order of Protein Secretion in Bacterial Type III Systems. Science (80-). 2011;331(6021):1188–91.

62. Jeckelmann JM, Erni B. Transporters of glucose and other carbohydrates in bacteria. Pflugers Arch Eur J Physiol. 2020;472(9):1129–53.

63. Pflüger-Grau K, Görke B. Regulatory roles of the bacterial nitrogen-related phosphotransferase system. Trends Microbiol. 2010;18(5):205–14.

64. Pacheco AR, Sperandio V. Enteric pathogens exploit the microbiota-generated nutritional environment of the gut. Metab Bact Pathog. 2015;(10):279–96.

65. Hong H, Patel DR, Tamm LK, van den Berg B. The Outer Membrane Protein OmpW Forms an Eight-stranded β-Barrel with a Hydrophobic Channel. J Biol Chem. 2006;281(11):7568–77.

66. Albrecht R, Zeth K, Söding J, Lupas A, Linke D. Expression, crystallization and preliminary X-ray crystallographic studies of the outer membrane protein OmpW from Escherichia coli. Acta Crystallogr Sect F Struct Biol Cryst Commun. 2006;62(4):415–8.

67. Wu X-B, Tian L-H, Zou H-J, Wang C-Y, Yu Z-Q, Tang C-H, et al. Outer membrane protein OmpW of Escherichia coli is required for resistance to phagocytosis. Res Microbiol. 2013;164(8):848–55.

68. Renault TT, Guse A, Erhardt M. Export Mechanisms and Energy Transduction in Type-III Secretion Machines. In: Current Topics in Microbiology and Immunology. Springer, Berlin, Heidelberg; 2019. p. 1–17.

69. Konovalova A, Silhavy TJ. Outer membrane lipoprotein biogenesis: Lol is not the end. Philos Trans R Soc B Biol Sci. 2015;370(1679).

70. Kudryashev M, Stenta M, Schmelz S, Amstutz M, Wiesand U, Castaño-Díez D, et al. In situ structural analysis of the Yersinia enterocolitica injectisome. Elife. 2013;2(Im):e00792.

71. Lambert deRouvroit C, Sluiters C, Cornelis GR. Role of the transcriptional activator, VirF, and temperature in the expression of the pYV plasmid genes of Yersinia enterocolitica. Mol Microbiol. 1992;6(3):395–409.

72. Schindelin J, Arganda-Carreras I, Frise E, Kaynig V, Longair M, Pietzsch T, et al. Fiji: an open-source platform for biological-image analysis. Nat Methods. 2012;9(7):676–82.

73. Thevenaz P, Ruttimann UE, Unser M. A pyramid approach to subpixel registration based on intensity. IEEE Trans Image Process. 1998;7(1):27–41.

74. Spahn C, Laine RF, Pereira P, Gómez-de-mariscal E. Supplementary information - DeepBacs: Bacterial image analysis using open-source deep learning approaches. bioRxiv. 2021;1–15.

75. Cox J, Mann M. MaxQuant enables high peptide identification rates, individualized p.p.b.-range mass accuracies and proteome-wide protein quantification. Nat Biotechnol. 2008;26(12):1367–72.

76. Fuchs TM, Brandt K, Starke M, Rattei T. Shotgun sequencing of Yersinia enterocolitica strain W22703 (biotype 2, serotype O:9): Genomic evidence for oscillation between invertebrates and mammals. BMC Genomics. 2011;12:1–14.

77. Glatter T, Ludwig C, Ahrné E, Aebersold R, Heck AJR, Schmidt A. Large-scale quantitative assessment of different in-solution protein digestion protocols reveals superior cleavage efficiency of tandem Lys-C/trypsin proteolysis over trypsin digestion. J Proteome Res. 2012;11(11):5145–56.

78. Ahrné E, Glatter T, Viganò C, Von Schubert C, Nigg EA, Schmidt A. Evaluation and improvement of quantification accuracy in isobaric mass tag-based protein quantification experiments. J Proteome Res. 2016;15(8):2537–47.

79. Tyanova S, Temu T, Sinitcyn P, Carlson A, Hein MY, Geiger T, et al. The Perseus computational platform for comprehensive analysis of (prote)omics data. Nat Methods. 2016;13(9):731–40.

80. Lampaki D, Diepold A, Glatter T. A Serial Sample Processing Strategy with Improved Performance for in-Depth Quantitative Analysis of Type III Secretion Events in Pseudomonas aeruginosa. J Proteome Res. 2020;19(1):543–53.

81. Wallace IM, O’Sullivan O, Higgins DG, Notredame C. M-Coffee: combining multiple sequence alignment methods with T-Coffee. Nucleic Acids Res. 2006;34:1692–9.

82. Erhardt M, Namba K, Hughes KT. Bacterial nanomachines: the flagellum and type III injectisome. Cold Spring Harb Perspect Biol. 2010;2(11):a000299.

83. Hueck CJ. Type III protein secretion systems in bacterial pathogens of animals and plants. Microbiol Mol Biol Rev. 1998;62(2):379–433.

84. Wagner S, Diepold A. A Unified Nomenclature for Injectisome-Type Type III Secretion Systems. In: Current topics in microbiology and immunology. 2020. p. 427: 1–10.

