## Supplementary Information for "Pilotins are mobile T3SS components involved in assembly and substrate specificity of the bacterial type III secretion system"

to

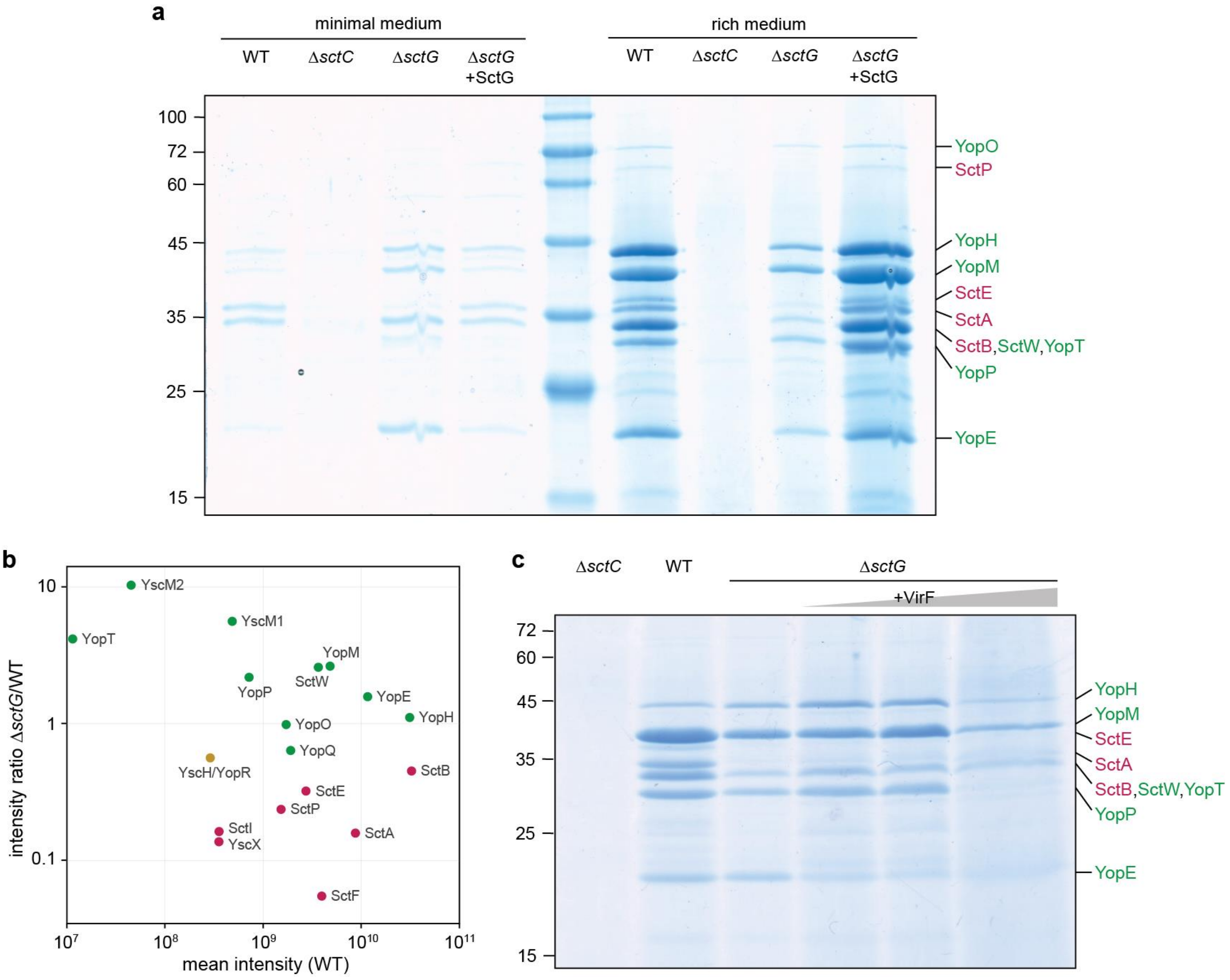

**Fig. S1: Complementation of protein secretion in strains lacking the pilotin SctG**

**a**) Secretion assay for the indicated strains in minimal medium (left) and in rich medium (BHI, right) as performed for the samples analyzed by mass spectrometry (Table 1). *In trans* complementation of SctG from plasmid induced by addition of 0.2% arabinose. Normalized amounts of supernatant (equivalent to $3 \times 10^8$ bacteria per lane) were loaded on a SDS-PAGE gel and stained with Coomassie Brilliant Blue. Left, molecular weight in kDa; right, assignment of exported proteins. *n*=5. **b**) Label-free mass spectrometry quantification of secreted proteins in Δ*sctG* and wild type (WT) in minimal medium, intensity ratio plotted against mean intensity (*n*=3). **c)** Secretion assay in rich medium for the indicated strains. Left, molecular weight in kDa. VirF expression from plasmid was induced with IPTG concentrations of 0, 10, 200 µM (from left to right). Overexpression of VirF by induction with 200 µM IPTG consistently decreased secretion and led to cell lysis, resulting in lower bacterial densities ($OD_{600}$ = 3.1±0.6, 3.0±0.6, 1.1±0.1 for 0, 10, 200 µM IPTG, respectively) and a widening of the respective lane on the gel. Left, molecular weight in kDa; right, assignment of exported proteins. *n*=3. In all panels, early/middle T3SS export substrates are marked in magenta; late substrates (effectors) are marked in green; substrate with unknown time point of export marked in yellow.

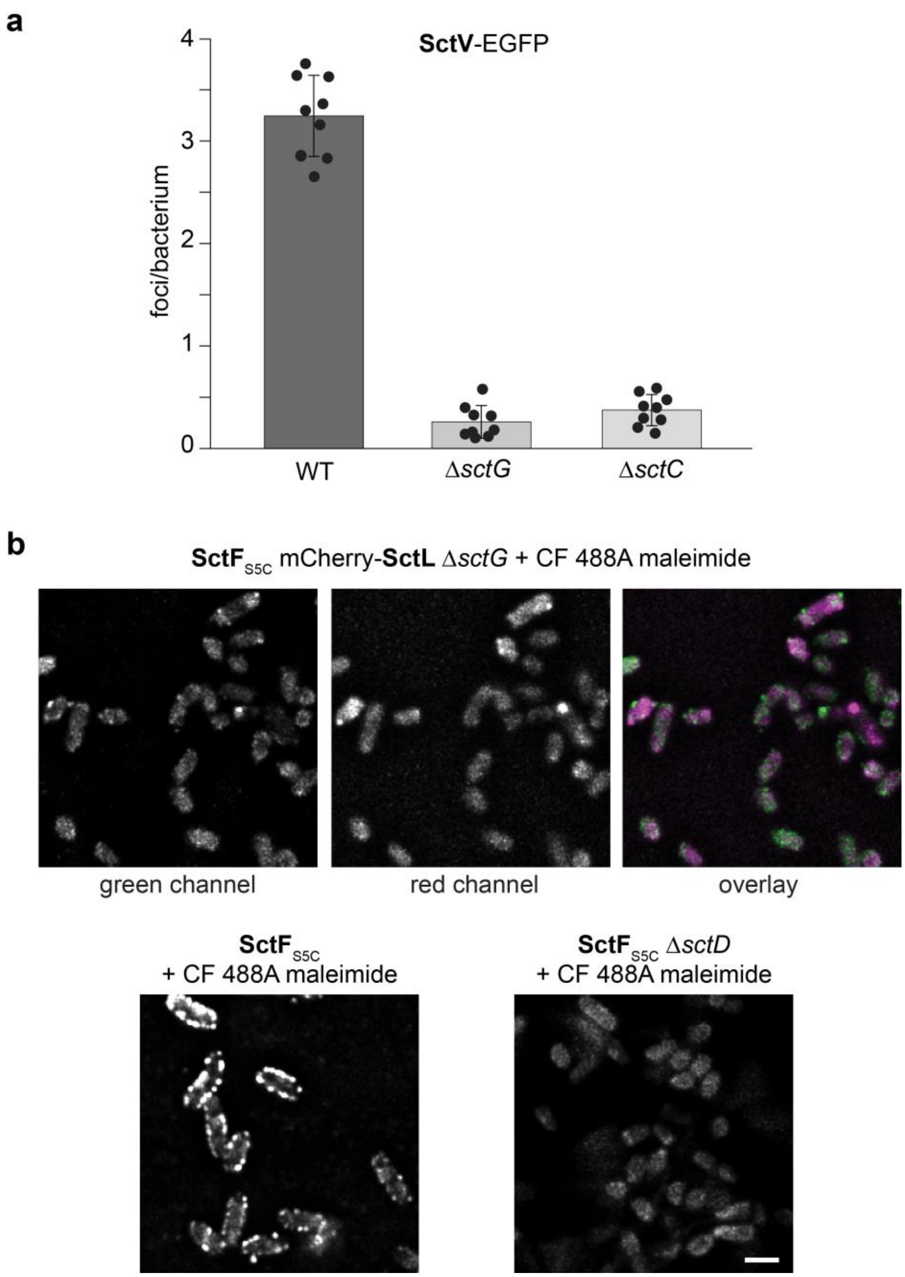

**Fig. S2: Absence of the pilotin leads to lack of stable anchoring of the export apparatus and formation of few needles and cytosolic sorting platforms, which do not colocalize**

**a**) Quantification of SctV-EGFP foci that are stable over a period of 90 s in the indicated strain backgrounds, as shown in Fig. 2b. Average detected foci per bacterium (see Material and Methods for details) in three fields of view from three independent experiments. Error bars and black circles indicate standard deviation and values of individual fields of view. **b**) Top, representative micrograph of $SctF_{S5C}$ stained with CF 488A maleimide (left and green in overlay) and mCherry-SctL (center and magenta in overlay) in a double labeled strain lacking SctG, as shown in Fig. 2a. Bottom, controls for needle staining; $SctF_{S5C}$ stained with CF 488A maleimide in strains otherwise wild-type (left) and lacking SctD (right). Scale bar, 2 µm. *n*=3.

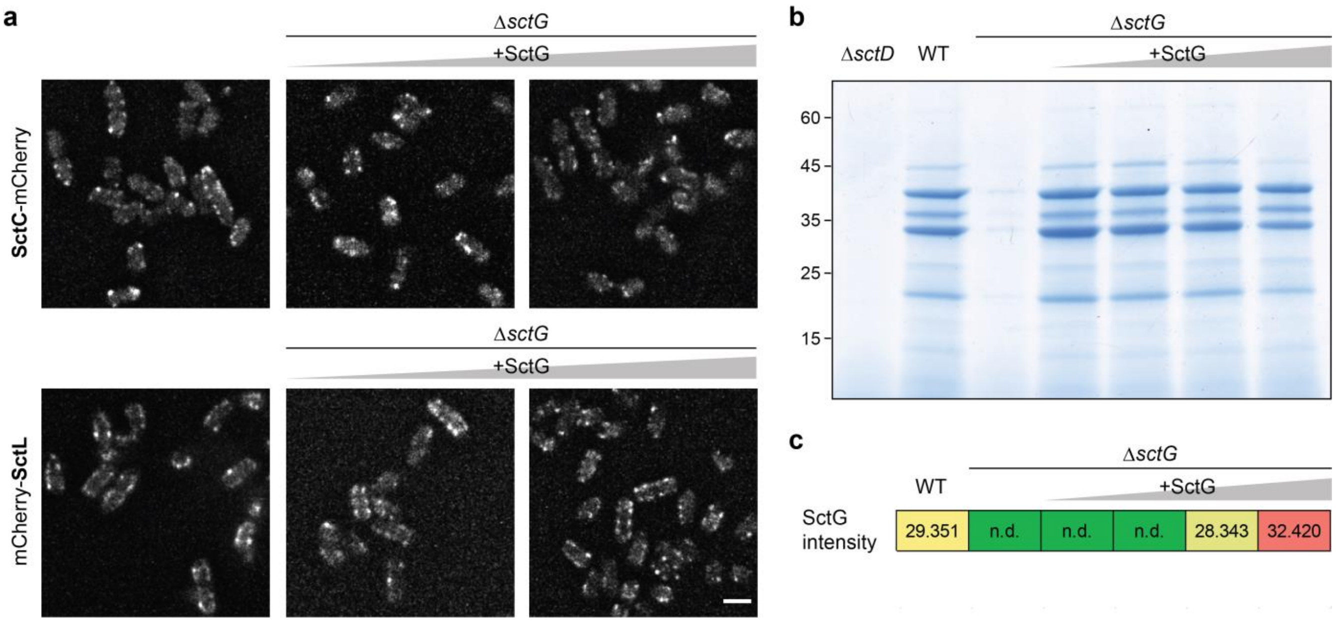

**Fig. S3: Lower than native amounts of SctG are sufficient to restore assembly of T3SS components and protein secretion**

**a**) Fluorescent micrographs of the indicated proteins in live *Y. enterocolitica.* Top, SctC-mCherry; bottom, mCherry-SctL in strains otherwise wild-type (left) and *ΔsctG*, complemented *in trans* by pBAD::SctG, induced with 0.02% or 0.2% L-arabinose (middle and right). Microscopy was performed under non-secreting conditions. Scale bar, 2 µm. *n*=3. **b**) Secretion assay showing the export of native T3SS substrates into the bacterial supernatant. Δ*sctD*, wild-type (WT), Δ*sctG* and Δ*sctG* complemented *in trans* by pBAD::SctG at increasing induction levels (from left to right: 0.2% glucose for repression; no addition; 0.02% arabinose; 0.2% arabinose). *n*=3. **c**) Label-free mass spectrometry quantification of SctG in total cellular protein samples of secreting *Y. enterocolitica* (from left to right: WT, Δ*sctG* and Δ*sctG* complemented *in trans* by pBAD::SctG at increasing induction levels as above) by label-free mass spectrometry (log 2 intensity values, average values of technical duplicate of one biological replicate). n.d., not detected.

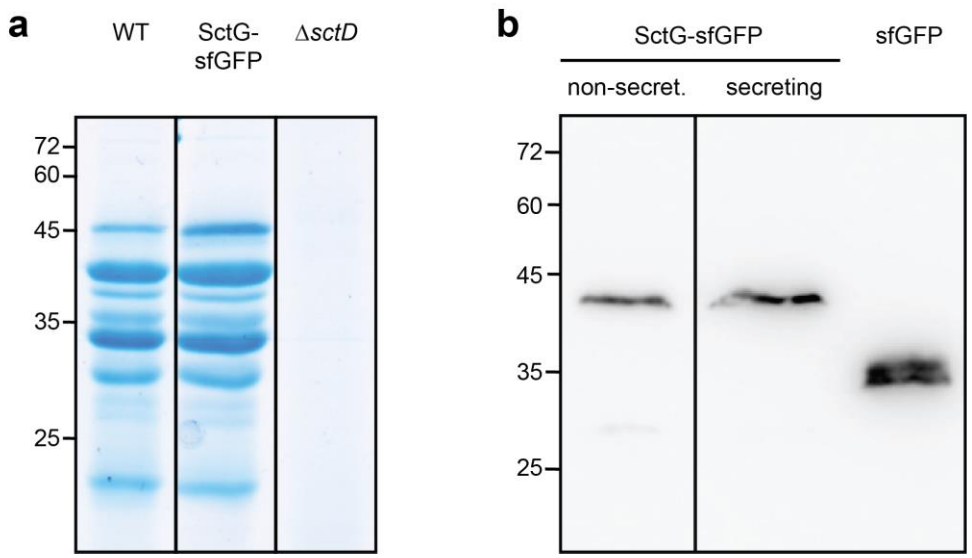

**Fig. S4: SctG can be functionally fluorescently labeled labelled at the C-terminus**

**a**) Secretion assay of the indicated strains showing the functionality of labeled SctG. Left, molecular weight in kDa. **b**) Stability of fusion proteins under non-secreting and secreting conditions. Immunoblot anti-GFP; left, molecular weight in kDa. Expected molecular weight of SctG-sfGFP: 42.2 kDa. sfGFP expressed from plasmid serves as a control. Vertical lines denote omission of intermediary lanes on the same gel.

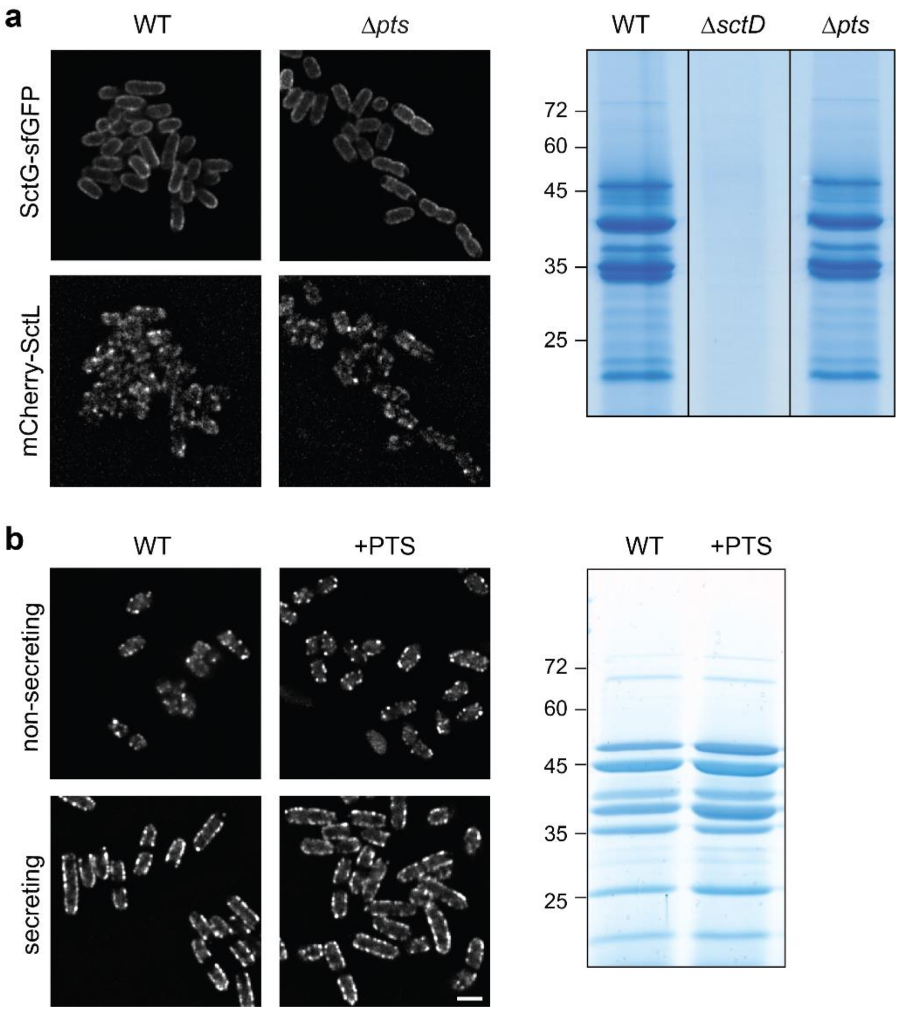

**Fig. S5 – The SctG interactor PTS trehalose transporter subunit EIIBC does not affect assembly or function of the T3SS in rich medium.**

**a**) Left: Micrographs of SctG-sfGFP and mCherry-SctL in double-labeled *Y. enterocolitica* strains otherwise wild-type (WT) or with a deletion of the PTS trehalose transporter subunit EIIBC (Δ*pts*). The experiment was performed under non-secreting conditions (*n*=3). Right: Secretion assay of the PTS trehalose transporter subunit EIIBC (Δ*pts*) deletion with WT and Δ*sctD* strains serving as positive and negative controls, under secreting conditions (*n*=2), molecular weight in kDa indicated. **b**) Left: Micrographs of EGFP-SctQ expressed from its native location on the virulence plasmid in *Y. enterocolitica* otherwise WT or overexpressing the PTS trehalose transporter subunit EIIBC (+PTS). Experiments performed under non-secreting and secreting conditions as indicated (*n*=3 each). Right: corresponding secretion assay, molecular weight in kDa indicated. Scale bar for all micrographs, 2 µm.

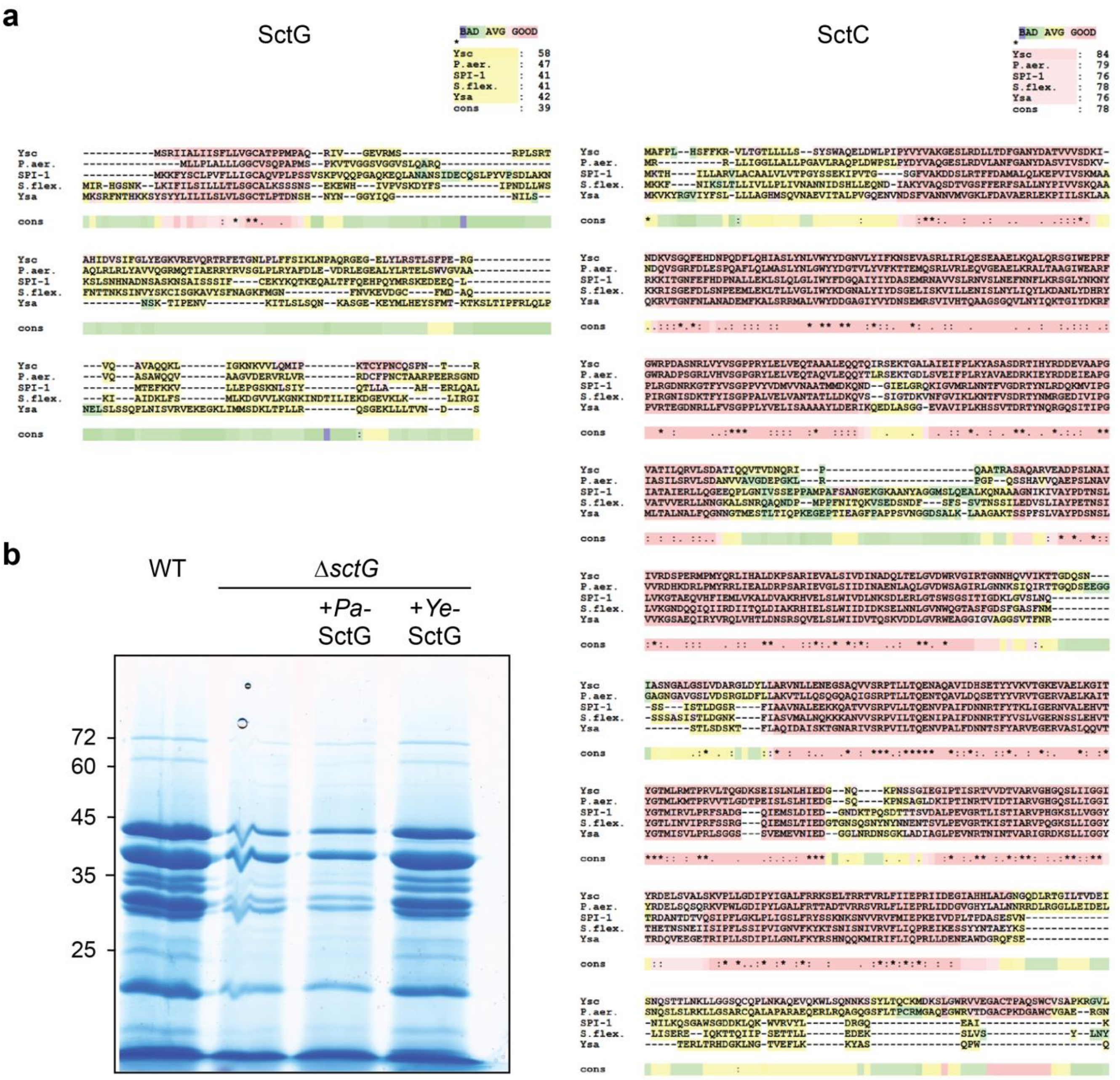

**Fig. S6: Limited sequence conservation and complementation of SctG homologs**

**a**) Multiple sequence alignments of SctG (left) and SctC (right) from different T3SS, assembled using T-COFFEE 11.0 [1]. Organisms and T3SS from top: *Y. enterocolitica* Ysc, *P. aeruginosa*, *Salmonella* Typhimurium SPI-1, *Shigella flexneri*, *Y. enterocolitica* Ysa. **b**) Secretion assay showing the export of native T3SS substrates in *Y. enterocolitica* strains with the indicated genotypes. A deletion of *sctG* can be complemented by *Y. enterocolitica* SctG (*Ye*SctG), but not the *P. aeruginosa* homolog (*Pa*SctG). Left, molecular weight in kDa.

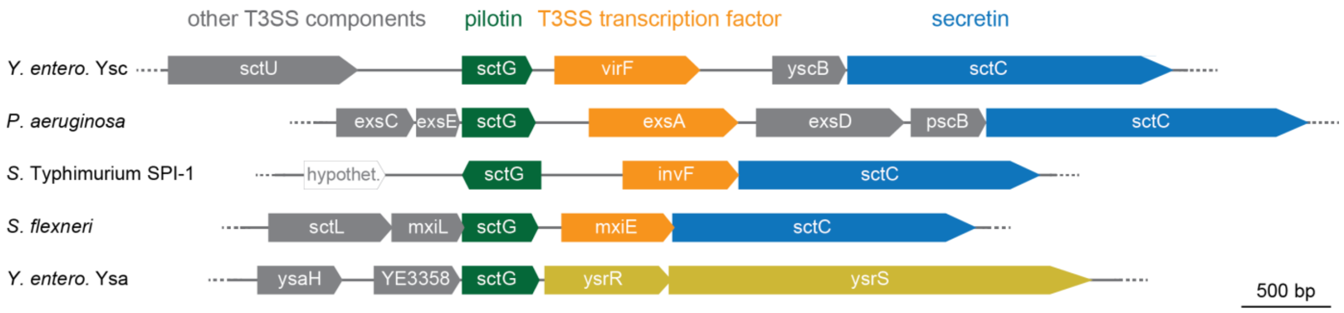

**Fig. S7 – Genetic neighborhood analysis of pilotin proteins**

Genetic neighborhood analysis of pilotin proteins (green) of the different T3SS indicated on the left side. Main T3SS transcription factors (orange), secretins (blue) and other T3SS components (gray) are indicated. The YsrRS phosphorelay system activating transcription of the Ysa system [2] is marked in yellow.

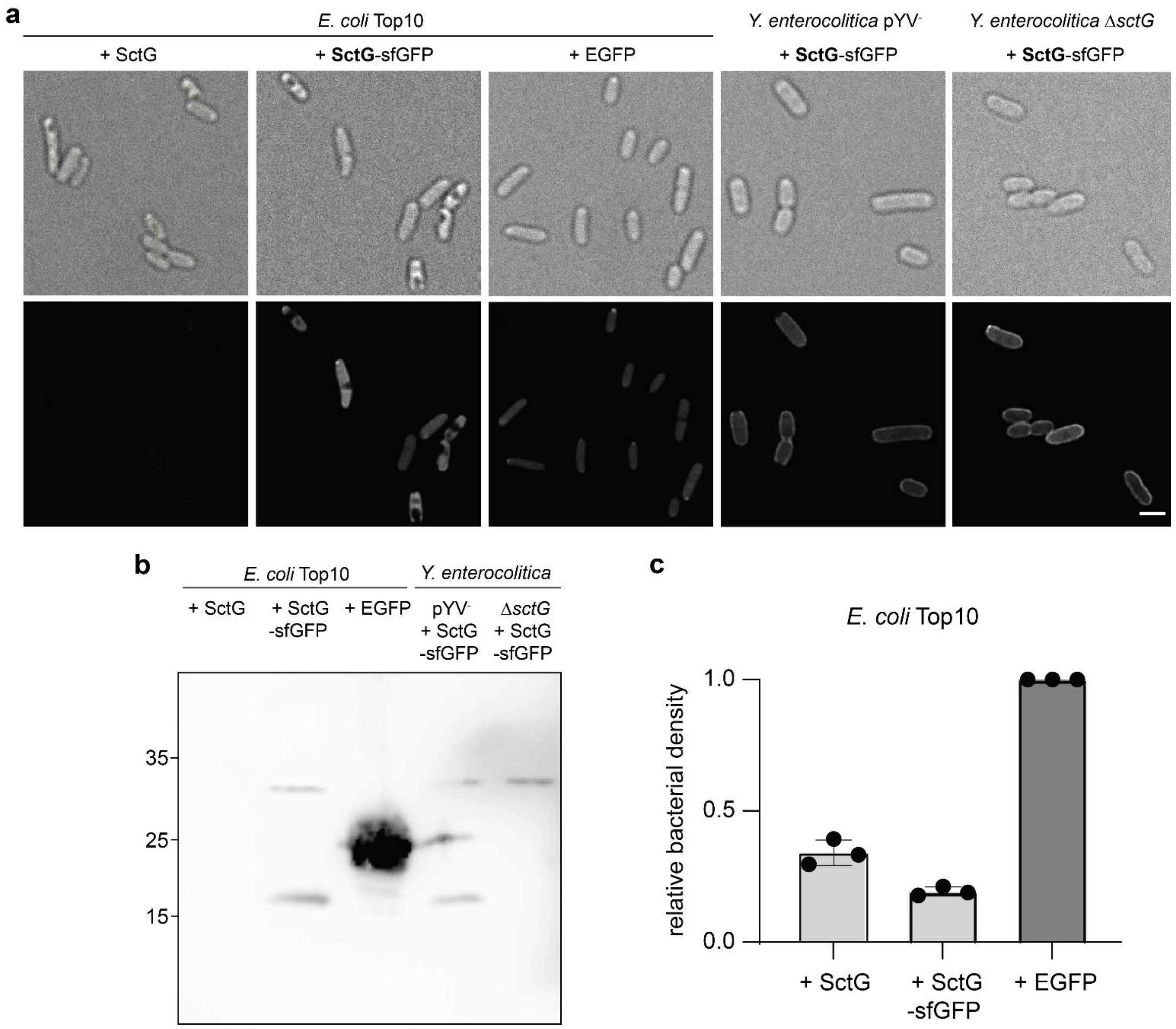

**Fig. S8: Localization of SctG-sfGFP and effect of pilotin expression on T3SS-negative bacteria.**

Ectopic expression of SctG and SctG-sfGFP from plasmid in *E. coli* Top10 and the indicated *Y. enterocolitica* mutants. In case of *Y. enterocolitica*, the experiment was performed under non-secreting conditions. Protein expression was induced with 0.02% arabinose. **a**) Light (top) and fluorescence (bottom) micrographs of the indicated proteins. **b**) Immunoblot of total cellular proteins with anti-GFP antibody. Left, molecular weight in kDa. **c**) Normalized $OD_{600}$ at the time of the experiment (after 3 hours of incubation at 37°C). *n*=3.

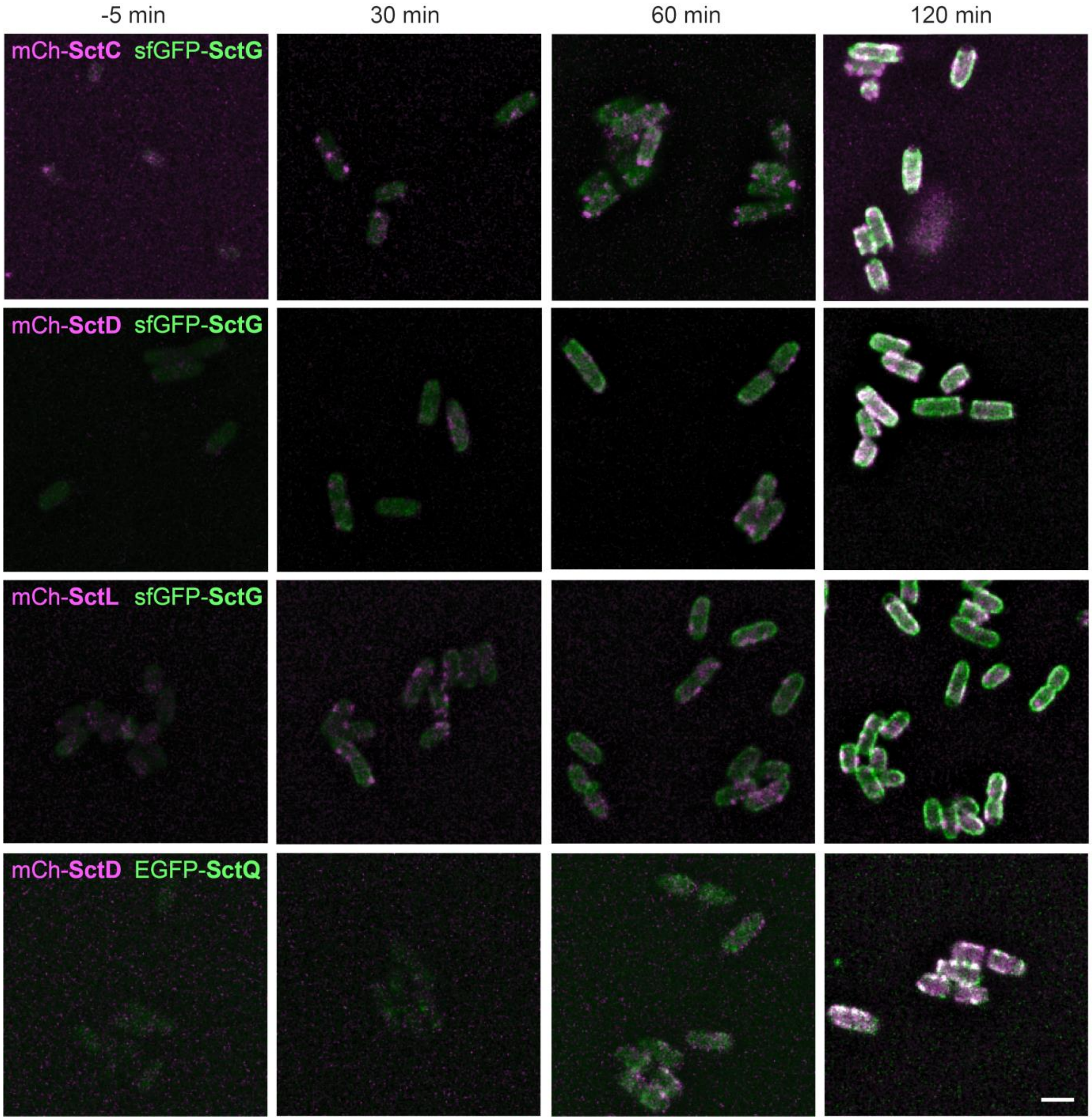

**Fig. S9 – SctG does not colocalize with any other injectisome component over time.**

Micrographs displaying a time course experiment with SctG-sfGFP (green) and different injectisome components labeled with mCherry (mCh-SctC/-SctD/SctL) (magenta) in the respective double-labeled strains under non-secreting conditions. Indicated times relative to shift to 37°C to induce expression of the *yop* regulon. A strain expressing mCherry-SctD and EGFP-SctQ was used as a control. *n*=3. Scale bar, 2 µm.

### Table S1 – Common and species-specific protein names for T3SS components

Common "Sct" nomenclature based on [3] with additions by [4,5]. Degree of conservation adapted from [6]. –, no clear homologue present; n.d., not determined (homologues present in less than three T3SS families).

| **Functional name** | **T3S family** | Ysc | Inv-Mxi-Spa | | Ssa-Esc | | Hrp-Hrc 1 | Hrp-Hrc 2 | **Degree of conservation (Alignment score)** | **Flagellar homo-logue** |
|---|---|---|---|---|---|---|---|---|---|---|
| | **Sct name** | ***Yersinia*** | ***Salmo-nella SPI-1*** | ***Shigella*** | ***Salmo-nella SPI-2*** | ***Esche-richia coli*** | ***Pseudo-monas syringae*** | ***Ralstonia solana-cearum*** | | |
| Secretin | **SctC** | YscC | InvG | MxiD | SsaC | EscC | HrcC | HrcC | high (75) | - |
| Outer MS ring protein | **SctD** | YscD | PrgH | MxiG | SsaD | EscD | HrpQ | HrpW | low (33) | FliG? |
| Inner MS ring protein | **SctJ** | YscJ | PrgK | MxiJ | SsaJ | EscJ | HrcJ | HrcJ | high (75) | FliF |
| Minor export apparatus protein | **SctR** | YscR | SpaP | Spa24 (SpaP) | SsaR | EscR | HrcR | HrcR | high (91) | FliP |
| Minor export apparatus protein | **SctS** | YscS | SpaQ | Spa9 (SpaQ) | SsaS | EscS | HrcS | HrcS | high (94) | FliQ |
| Minor export apparatus protein | **SctT** | YscT | SpaR | Spa29 (SpaR) | SsaT | EscT | HrcT | HrcT | high (83) | FliR |
| Export apparatus switch protein | **SctU** | YscU | SpaS | Spa40 (SpaS) | SsaU | EscU | HrcU | HrcU | high (84) | FlhB |
| Major export apparatus protein | **SctV** | YscV | InvA | MxiA | SsaV | EscV | HrcV | HrcV | high (78) | FlhA |
| Accessory cytosolic protein | **SctK** | YscK | OrgA | MxiK | - | - | HrpD | - | low (38) | - |
| C ring/pod protein | **SctQ** | YscQ | SpaO | Spa33 (SpaO) | SsaQ | EscQ | $HrcQ_{A+B}$ | HrcQ | low (38) | FliM+FliN |
| Stator | **SctL** | YscL | OrgB | MxiN | SsaK | EscL (Orf5) | HrpE | HrpF | low (45) | FliH |
| ATPase | **SctN** | YscN | InvC | Spa47 (SpaL) | SsaN | EscN | HrcN | HrcN | high (92) | FliI |
| Stalk | **SctO** | YscO | InvI | Spa13 (SpaM) | SsaO | Orf15 | HrpO | HrpD | very low (27) | FliJ |
| Needle filament protein | **SctF** | YscF | PrgI | MxiH | SsaG | EscF | HrpA | HrpY | low (40) | - |
| Inner rod / washer protein | **SctI** | YscI | PrgJ | MxiI | SsaI | EscI (rOrf8) | HrpB | HrpJ | low (47) | - |
| Needle length regulator | **SctP** | YscP | InvJ | Spa32 (SpaN) | SsaP | EscP (Orf16) | HrpP | HpaP | low (33) | FliK |
| Hydrophilic translocator, needle tip protein | **SctA** | LcrV | SipD | IpaD | - | EspA? | - | - | n.d. | - |
| Hydrophobic translocator, pore protein | **SctE** | YopB | SipB | IpaB | SseC | EspD | HrpK | PopF1, PopF2 | very low (21) | - |
| Hydrophobic translocator, pore protein | **SctB** | YopD | SipC | IpaC | SseD | EspB | - | - | very low (27) | - |
| **Pilotin** | SctG | **YscW** | **InvH** | **MxiM** | **-** | **-** | **-** | **-** | **n.d.** | **-** |
| Gatekeeper | **SctW** | YopN | InvE | MxiC | SsaL | SepL | HrpJ | HpaA | low (36) | - |

**Table S2 - SctG has an influence on the expression of T3SS components, but very few other proteins**

Comparison of expression levels in total cellular protein samples of wild-type (WT) and *ΔsctG* (dSctG) *Y. enterocolitica* under non-secreting conditions, determined by label-free mass spectrometry.

All proteins with $p < 0.01$ and number of detected peptides (# pept.) > 2 are included in the list, and ordered inversely according to their expression ratio *ΔsctG* / WT. SctG is shaded in orange, other T3SS components are shaded in yellow, except for the main transcriptional activator VirF, which is shaded in blue. Non-T3SS interactors discussed in the text are shaded in green. The PTS trehalose export subunit II BC and exported T3SS proteins included in Table 1 are additionally listed at the end of the table for added information, see main text for details.

| Protein | Mean intensity | | log 2 intensity ratio | *p* value | Individual intensities | | | | | | # pept. |
|---|---|---|---|---|---|---|---|---|---|---|---|
| | WT | dSctG | | | WT 1 | WT 2 | WT 3 | dSctG 1 | dSctG 2 | dSctG 3 | |
| T3SS lipoprotein component SctG (YscW) | 2.24E+08 | 7.03E+06 | -6.493 | 0.0085 | 2.1E+08 | 2.3E+08 | 2.3E+08 | 1.3E+07 | 8.1E+06 | 1.5E+05 | 9 |
| Outer membrane protein W | 1.61E+08 | 2.65E+07 | -2.913 | 0.0078 | 2.4E+08 | 1.1E+08 | 1.3E+08 | 4.5E+07 | 6.7E+06 | 2.7E+07 | 3 |
| T3SS component SctD | 8.28E+07 | 1.72E+07 | -2.274 | 0.0000 | 9.2E+07 | 7.3E+07 | 8.3E+07 | 2.0E+07 | 1.7E+07 | 1.5E+07 | 13 |
| 5-methyltetrahydropteroyltriglutamate-homocysteine methyltransferase | 3.44E+07 | 9.99E+06 | -1.842 | 0.0015 | 2.7E+07 | 3.8E+07 | 3.9E+07 | 9.5E+06 | 6.3E+06 | 1.4E+07 | 6 |
| Succinate dehydrogenase iron-sulfur subunit | 2.34E+07 | 8.69E+06 | -1.502 | 0.0022 | 2.3E+07 | 2.3E+07 | 2.4E+07 | 6.0E+06 | 1.3E+07 | 7.3E+06 | 3 |
| T3SS transcriptional activator VirF | 4.58E+07 | 1.66E+07 | -1.451 | 0.0004 | 5.1E+07 | 5.1E+07 | 3.5E+07 | 1.8E+07 | 1.5E+07 | 1.7E+07 | 4 |
| T3SS effector YopQ | 9.54E+07 | 3.84E+07 | -1.330 | 0.0008 | 1.1E+08 | 9.4E+07 | 8.5E+07 | 3.4E+07 | 4.9E+07 | 3.2E+07 | 12 |
| T3SS component SctF | 3.76E+08 | 1.58E+08 | -1.288 | 0.0011 | 3.7E+08 | 3.9E+08 | 3.6E+08 | 1.2E+08 | 1.6E+08 | 2.0E+08 | 4 |
| T3SS effector YopO | 4.72E+07 | 2.06E+07 | -1.286 | 0.0064 | 5.0E+07 | 4.5E+07 | 4.7E+07 | 1.5E+07 | 3.1E+07 | 1.6E+07 | 12 |
| T3SS protein SctB | 4.84E+08 | 2.17E+08 | -1.220 | 0.0062 | 4.4E+08 | 5.3E+08 | 4.8E+08 | 1.3E+08 | 2.3E+08 | 2.8E+08 | 4 |
| T3SS component SctI | 3.15E+07 | 1.34E+07 | -1.215 | 0.0005 | 3.1E+07 | 2.6E+07 | 3.7E+07 | 1.4E+07 | 1.4E+07 | 1.3E+07 | 5 |
| T3SS effector effector YopH | 6.08E+08 | 2.67E+08 | -1.199 | 0.0003 | 6.1E+08 | 6.1E+08 | 6.0E+08 | 2.6E+08 | 2.3E+08 | 3.1E+08 | 32 |
| Hemin receptor | 5.71E+08 | 2.51E+08 | -1.192 | 0.0007 | 6.5E+08 | 5.3E+08 | 5.4E+08 | 2.0E+08 | 2.7E+08 | 2.8E+08 | 18 |
| Putative uncharacterized protein | 3.92E+08 | 1.80E+08 | -1.129 | 0.0004 | 4.3E+08 | 3.7E+08 | 3.7E+08 | 1.5E+08 | 1.8E+08 | 2.0E+08 | 5 |
| Anaerobic glycerol-3-phosphate dehydrogenase subunit B | 2.79E+07 | 1.34E+07 | -1.121 | 0.0069 | 3.0E+07 | 2.6E+07 | 2.8E+07 | 1.1E+07 | 1.0E+07 | 1.9E+07 | 6 |
| 3D-(3,5/4)-trihydroxycyclohexane-1,2-dionehydrolase | 2.10E+07 | 1.00E+07 | -1.089 | 0.0077 | 1.7E+07 | 2.6E+07 | 2.0E+07 | 1.3E+07 | 1.0E+07 | 7.1E+06 | 6 |
| T3SS effector YopE | 1.28E+09 | 6.78E+08 | -0.939 | 0.0025 | 1.3E+09 | 1.3E+09 | 1.3E+09 | 5.3E+08 | 6.9E+08 | 8.1E+08 | 9 |

| | | | | | | | | | | | |
|---|---|---|---|---|---|---|---|---|---|---|---|
| T3SS chaperone SycN | 4.43E+08 | 2.36E+08 | -0.928 | 0.0028 | 4.6E+08 | 4.6E+08 | 4.1E+08 | 1.8E+08 | 2.5E+08 | 2.7E+08 | 5 |
| T3SS chaperone YscG | 4.23E+08 | 2.27E+08 | -0.922 | 0.0068 | 4.4E+08 | 4.5E+08 | 3.7E+08 | 1.7E+08 | 2.8E+08 | 2.3E+08 | 5 |
| T3SS translocator SctE (YopB) | 3.08E+08 | 1.66E+08 | -0.896 | 0.0006 | 3.3E+08 | 3.0E+08 | 2.9E+08 | 1.6E+08 | 1.8E+08 | 1.6E+08 | 15 |
| Taurine-binding periplasmic protein | 9.63E+07 | 5.28E+07 | -0.873 | 0.0011 | 9.8E+07 | 1.0E+08 | 9.1E+07 | 4.7E+07 | 5.1E+07 | 6.0E+07 | 11 |
| T3SS gatekeeper SctW (YopN) | 2.24E+08 | 1.23E+08 | -0.870 | 0.0008 | 2.1E+08 | 2.4E+08 | 2.2E+08 | 1.3E+08 | 1.1E+08 | 1.3E+08 | 5 |
| T3SS chaperone YscE | 4.23E+08 | 2.39E+08 | -0.840 | 0.0049 | 4.4E+08 | 4.4E+08 | 3.9E+08 | 1.9E+08 | 2.8E+08 | 2.5E+08 | 7 |
| Putative uncharacterized protein (Fragment) | 1.41E+07 | 8.07E+06 | -0.817 | 0.0037 | 1.4E+07 | 1.4E+07 | 1.3E+07 | 9.0E+06 | 8.8E+06 | 6.4E+06 | 3 |
| T3SS translocator SctB (YopD) | 1.11E+09 | 6.51E+08 | -0.799 | 0.0090 | 1.0E+09 | 1.2E+09 | 1.1E+09 | 5.2E+08 | 5.9E+08 | 8.4E+08 | 14 |
| Hemin transport protein HemS | 8.93E+08 | 5.27E+08 | -0.775 | 0.0041 | 9.5E+08 | 8.6E+08 | 8.8E+08 | 4.4E+08 | 5.3E+08 | 6.2E+08 | 20 |
| T3SS component SctN | 1.06E+08 | 6.24E+07 | -0.764 | 0.0011 | 1.1E+08 | 1.1E+08 | 1.0E+08 | 6.1E+07 | 6.0E+07 | 6.6E+07 | 17 |
| DNA replication terminus site-binding protein | 2.38E+07 | 1.42E+07 | -0.756 | 0.0027 | 2.4E+07 | 2.4E+07 | 2.3E+07 | 1.2E+07 | 1.6E+07 | 1.4E+07 | 4 |
| Nitrite reductase [NAD(P)H] large subunit | 3.29E+07 | 1.96E+07 | -0.737 | 0.0069 | 3.7E+07 | 2.7E+07 | 3.5E+07 | 2.0E+07 | 1.8E+07 | 2.1E+07 | 13 |
| T3SS chaperone SycD | 1.66E+08 | 1.01E+08 | -0.729 | 0.0077 | 1.8E+08 | 1.6E+08 | 1.6E+08 | 8.0E+07 | 1.1E+08 | 1.2E+08 | 6 |
| T3SS chaperone SycE | 1.29E+09 | 7.96E+08 | -0.706 | 0.0045 | 1.4E+09 | 1.3E+09 | 1.2E+09 | 6.7E+08 | 8.3E+08 | 8.8E+08 | 5 |
| 6-carboxy-5,6,7,8-tetrahydropterin synthase | 9.53E+07 | 5.92E+07 | -0.706 | 0.0080 | 9.6E+07 | 9.4E+07 | 9.5E+07 | 4.9E+07 | 7.2E+07 | 5.7E+07 | 4 |
| T3SS component SctJ | 2.72E+08 | 1.69E+08 | -0.699 | 0.0055 | 2.7E+08 | 2.9E+08 | 2.6E+08 | 1.4E+08 | 2.0E+08 | 1.7E+08 | 13 |
| T3SS component SctL | 3.28E+08 | 2.05E+08 | -0.694 | 0.0084 | 3.1E+08 | 3.3E+08 | 3.5E+08 | 1.6E+08 | 2.2E+08 | 2.4E+08 | 9 |
| T3SS component SctQ | 1.29E+09 | 9.12E+08 | -0.507 | 0.0098 | 1.3E+09 | 1.3E+09 | 1.2E+09 | 8.4E+08 | 9.0E+08 | 9.9E+08 | 10 |
| Succinate dehydrogenase flavoprotein subunit | 3.84E+08 | 5.74E+08 | 0.580 | 0.0046 | 3.7E+08 | 4.1E+08 | 3.8E+08 | 5.4E+08 | 5.7E+08 | 6.1E+08 | 25 |
| Cysteine desulfurase IscS | 5.73E+08 | 8.58E+08 | 0.586 | 0.0066 | 5.1E+08 | 6.1E+08 | 6.0E+08 | 8.6E+08 | 8.1E+08 | 9.0E+08 | 13 |
| Phage shock protein B | 7.04E+07 | 1.28E+08 | 0.855 | 0.0019 | 7.0E+07 | 7.7E+07 | 6.5E+07 | 1.1E+08 | 1.2E+08 | 1.5E+08 | 3 |
| Phage shock protein A | 1.74E+08 | 3.81E+08 | 1.110 | 0.0021 | 1.5E+08 | 1.9E+08 | 1.8E+08 | 2.9E+08 | 3.8E+08 | 4.7E+08 | 17 |
| HTH-type transcriptional regulator IscR | 8.15E+07 | 1.83E+08 | 1.114 | 0.0071 | 7.3E+07 | 8.9E+07 | 8.3E+07 | 1.3E+08 | 2.5E+08 | 1.7E+08 | 5 |
| Putative uncharacterized protein | 5.96E+06 | 6.22E+07 | 3.393 | 0.0000 | 5.2E+06 | 7.5E+06 | 5.3E+06 | 7.3E+07 | 5.8E+07 | 5.6E+07 | 3 |
| *PTS trehalose transporter subunit EIIBC* | *5.30E+07* | *7.11E+07* | *-0.049* | *0.9699* | *6.2E+07* | *1.3E+07* | *8.3E+07* | *1.5E+08* | *6.1E+07* | *7.1E+06* | *7* |
| T3SS effector YopM | 3.86E+08 | 2.51E+08 | -0.648 | 0.0201 | 3.8E+08 | 3.9E+08 | 3.8E+08 | 2.3E+08 | 2.0E+08 | 3.2E+08 | 3 |
| T3SS regulator YscM2 | 2.72E+08 | 1.85E+08 | -0.631 | 0.1017 | 3.1E+08 | 2.6E+08 | 2.4E+08 | 1.1E+08 | 2.2E+08 | 2.3E+08 | 6 |

| | | | | | | | | | | | |
|---|---|---|---|---|---|---|---|---|---|---|---|
| T3SS regulator YscM1 | 4.28E+08 | 3.03E+08 | -0.501 | 0.0170 | 4.4E+08 | 4.6E+08 | 3.8E+08 | 2.7E+08 | 3.2E+08 | 3.2E+08 | 8 |
| T3SS early substrate / regulator YscX | 6.45E+07 | 5.01E+07 | -0.382 | 0.0829 | 6.5E+07 | 6.6E+07 | 6.2E+07 | 4.2E+07 | 6.2E+07 | 4.7E+07 | 3 |
| T3SS needle length regulator / ruler SctP | 1.50E+08 | 1.24E+08 | -0.284 | 0.1529 | 1.5E+08 | 1.5E+08 | 1.5E+08 | 1.0E+08 | 1.3E+08 | 1.5E+08 | 16 |
| T3SS translocator SctA (LcrV) | 3.73E+08 | 3.44E+08 | -0.119 | 0.4137 | 3.6E+08 | 3.9E+08 | 3.6E+08 | 3.2E+08 | 3.4E+08 | 3.7E+08 | 16 |
| T3SS secreted protein YscH/YopR | 2.26E+07 | 1.79E+07 | 0.128 | 0.8911 | 8.8E+06 | 5.3E+07 | 6.3E+06 | 3.2E+07 | 1.2E+07 | 9.8E+06 | 2 |

**Table S3 – Intensities of individual experiments for Table 1**

Label-free quantitative mass spectrometry of exported known T3SS substrates in different strain backgrounds, as shown in Table 1. Top, measurement in rich medium (BHI); bottom, in minimal medium (see material and methods for details). Hits sorted by ratio Δ*sctG* / wild-type (WT) in mean intensities. Light grey font, imputed values (no peptides detected). All exported proteins with number of detected peptides (# pept.) > 2 are included in the list, which covers all expected secreted proteins except for SctO, for which only one peptide was detected.

| ***Rich medium*** | **Single intensities** | | | | | | | | | |
|---|---|---|---|---|---|---|---|---|---|---|
| **T3SS export substrate** | **WT 1** | **WT 2** | **WT 3** | **Δ*sctG* 1** | **Δ*sctG* 2** | **Δ*sctG* 3** | **Δ*sctD* 1** | **Δ*sctD* 2** | **Δ*sctD* 3** | **# pept.** |
| Translocator SctE (YopB) | 5.3E+08 | 3.6E+08 | 2.7E+08 | 2.4E+06 | 3.2E+06 | 1.3E+06 | 2.2E+05 | 1.4E+04 | 7.5E+05 | 29 |
| Translocator SctB (YopD) | 1.3E+10 | 1.0E+10 | 3.2E+09 | 1.3E+08 | 2.3E+08 | 1.0E+08 | 2.5E+06 | 3.6E+06 | 2.1E+06 | 73 |
| Tip protein SctA (LcrV) | 9.1E+08 | 8.7E+08 | 1.6E+08 | 1.1E+07 | 1.7E+07 | 8.3E+06 | 7.7E+04 | 1.0E+05 | 6.2E+04 | 41 |
| Needle component SctF | 3.5E+08 | 3.3E+08 | 5.7E+07 | 6.9E+06 | 7.8E+06 | 4.7E+06 | 1.2E+05 | 9.5E+04 | 2.7E+05 | 8 |
| Early substrate / regulator YscX | 7.9E+07 | 8.4E+07 | 1.9E+07 | 2.1E+06 | 2.5E+06 | 1.2E+06 | 5.0E+05 | 4.6E+04 | 3.7E+05 | 9 |
| Secreted protein YscH/YopR | 1.5E+08 | 1.4E+08 | 2.2E+07 | 3.4E+06 | 4.4E+06 | 2.5E+06 | 3.7E+04 | 5.3E+04 | 6.5E+04 | 8 |
| Rod / washer protein SctI | 1.7E+08 | 1.5E+08 | 3.8E+07 | 5.8E+06 | 7.3E+06 | 3.9E+06 | 8.9E+04 | 2.7E+04 | 4.1E+04 | 8 |
| Needle length regulator / ruler SctP | 3.2E+08 | 2.9E+08 | 1.2E+08 | 1.6E+07 | 2.5E+07 | 1.2E+07 | 1.1E+06 | 4.3E+05 | 8.5E+05 | 33 |
| Effector YopP | 2.2E+08 | 1.8E+08 | 1.0E+08 | 1.3E+07 | 2.2E+07 | 1.0E+07 | 1.3E+05 | 5.8E+04 | 4.0E+04 | 41 |
| Effector YopQ | 9.5E+08 | 8.1E+08 | 2.6E+08 | 5.4E+07 | 8.4E+07 | 4.2E+07 | 1.3E+06 | 8.5E+05 | 3.2E+05 | 24 |
| Effector YopO | 5.5E+08 | 3.9E+08 | 2.2E+08 | 4.3E+07 | 5.7E+07 | 3.7E+07 | 4.7E+04 | 2.9E+05 | 2.7E+04 | 71 |
| Effector YopT | 3.8E+07 | 2.5E+07 | 1.6E+07 | 3.2E+06 | 4.1E+06 | 2.5E+06 | 3.6E+04 | 2.3E+04 | 6.9E+04 | 13 |
| Effector YopE | 5.2E+09 | 3.7E+09 | 1.6E+09 | 4.8E+08 | 7.2E+08 | 3.4E+08 | 9.2E+05 | 8.7E+05 | 5.6E+05 | 25 |
| Gatekeeper SctW (YopN) | 2.5E+09 | 2.4E+09 | 4.7E+08 | 3.4E+08 | 4.7E+08 | 2.2E+08 | 1.2E+06 | 1.0E+06 | 9.6E+05 | 19 |
| Effector YopH | 2.4E+10 | 2.3E+10 | 6.1E+09 | 3.1E+09 | 4.9E+09 | 2.2E+09 | 3.2E+06 | 3.2E+06 | 3.1E+06 | 122 |
| Effector YopM | 3.6E+09 | 3.0E+09 | 6.1E+08 | 5.5E+08 | 7.9E+08 | 4.1E+08 | 2.0E+06 | 1.2E+06 | 1.2E+06 | 8 |
| Regulator YscM2 | 2.9E+07 | 3.1E+07 | 5.5E+06 | 6.3E+06 | 9.1E+06 | 4.5E+06 | 2.9E+04 | 7.0E+04 | 7.2E+04 | 9 |
| Regulator YscM1 | 1.2E+09 | 1.2E+09 | 2.0E+08 | 3.0E+08 | 4.0E+08 | 2.1E+08 | 1.0E+06 | 6.5E+05 | 6.2E+05 | 21 |

| ***Minimal medium*** | **Single intensities** | | | | | | | | | |
|---|---|---|---|---|---|---|---|---|---|---|
| **T3SS export substrate** | **WT 1** | **WT 2** | **WT 3** | **Δ*sctG* 1** | **Δ*sctG* 2** | **Δ*sctG* 3** | **Δ*sctC* 1** | **Δ*sctC* 2** | **Δ*sctC* 3** | **# pept.** |
| Needle component SctF | 3.3E+09 | 3.6E+09 | 4.9E+09 | 4.2E+08 | 1.9E+08 | 4.1E+07 | 2.4E+07 | 1.6E+07 | 9.2E+06 | 7 |
| Early substrate / regulator YscX | 2.0E+08 | 4.7E+08 | 4.0E+08 | 7.7E+07 | 2.1E+07 | 4.7E+07 | 1.2E+07 | 7.6E+06 | 6.7E+06 | 4 |
| Tip protein SctA (LcrV) | 8.1E+09 | 1.1E+10 | 7.5E+09 | 3.6E+09 | 4.6E+08 | 9.8E+07 | 1.1E+08 | 1.7E+08 | 4.0E+07 | 24 |
| Rod / washer protein SctI | 3.6E+08 | 3.9E+08 | 3.2E+08 | 8.9E+07 | 3.0E+07 | 5.4E+07 | 4.0E+06 | 5.4E+07 | 4.2E+06 | 3 |
| Needle length regulator / ruler SctP | 1.2E+09 | 2.1E+09 | 1.3E+09 | 6.8E+08 | 3.1E+08 | 9.1E+07 | 1.2E+07 | 1.3E+08 | 8.0E+06 | 23 |
| Translocator SctE (YopB) | 2.9E+09 | 3.8E+09 | 1.5E+09 | 2.1E+09 | 2.9E+08 | 2.7E+08 | 2.1E+07 | 2.6E+08 | 7.5E+08 | 19 |
| Translocator SctB (YopD) | 2.3E+10 | 4.4E+10 | 3.0E+10 | 3.6E+10 | 3.9E+09 | 3.7E+09 | 4.2E+08 | 2.0E+09 | 2.5E+08 | 46 |
| Secreted protein YscH/YopR | 3.0E+08 | 2.2E+08 | 3.5E+08 | 1.5E+08 | 3.9E+07 | 3.0E+08 | 1.0E+07 | 1.0E+07 | 1.4E+07 | 4 |
| Effector YopQ | 1.2E+09 | 2.8E+09 | 1.7E+09 | 2.4E+09 | 3.8E+08 | 8.7E+08 | 6.9E+06 | 7.3E+06 | 5.3E+06 | 16 |
| Effector YopO | 1.0E+09 | 2.1E+09 | 2.0E+09 | 3.6E+09 | 6.6E+08 | 7.6E+08 | 5.4E+06 | 1.8E+08 | 9.7E+06 | 37 |
| Effector YopH | 3.0E+10 | 3.3E+10 | 3.0E+10 | 4.9E+10 | 4.2E+10 | 1.2E+10 | 7.3E+08 | 2.4E+09 | 6.0E+08 | 64 |
| Effector YopE | 7.8E+09 | 1.5E+10 | 1.2E+10 | 3.3E+10 | 1.1E+10 | 1.1E+10 | 5.4E+08 | 7.3E+09 | 1.3E+08 | 10 |
| Effector YopP | 3.7E+08 | 1.1E+09 | 6.9E+08 | 4.2E+09 | 2.8E+08 | 2.8E+08 | 7.6E+06 | 3.8E+06 | 4.4E+06 | 25 |
| Gatekeeper SctW (YopN) | 2.7E+09 | 3.9E+09 | 4.4E+09 | 1.9E+10 | 2.4E+09 | 7.4E+09 | 2.7E+07 | 4.8E+07 | 2.1E+07 | 17 |
| Effector YopM | 4.9E+09 | 2.8E+09 | 6.8E+09 | 1.7E+10 | 4.2E+09 | 1.7E+10 | 6.7E+07 | 1.2E+08 | 5.2E+07 | 6 |
| Effector YopT | 1.3E+07 | 1.9E+07 | 2.5E+06 | 1.3E+08 | 8.8E+06 | 7.3E+06 | 6.0E+06 | 2.1E+07 | 1.2E+07 | 7 |
| Regulator YscM1 | 5.3E+08 | 3.8E+08 | 5.4E+08 | 3.9E+09 | 2.6E+09 | 1.7E+09 | 8.2E+06 | 1.4E+07 | 7.3E+06 | 12 |
| Regulator YscM2 | 8.1E+06 | 5.7E+07 | 7.1E+07 | 8.4E+08 | 3.2E+08 | 2.3E+08 | 6.4E+06 | 6.5E+06 | 3.4E+06 | 7 |

**Table S4 - SctG can be functionally labelled at the C-terminus**

Comparison of expression levels in total cellular protein samples of wild-type (WT) and SctG-sfGFP *Y. enterocolitica* under non-secreting conditions, determined by label-free mass spectrometry.

All proteins with $p < 0.01$ and number of detected peptides (# pept.) > 2 are included in the list, and ordered according to their expression ratio SctG-sfGFP / wild-type (WT). SctG is shaded in orange, other T3SS components are shaded in yellow. The T3SS main transcriptional regulator VirF (shaded in blue) is included at the bottom of the list for added information.

| Protein | Mean intensity | | log 2 intensity ratio | *p* value | Single intensities | | | | | | # pept. |
|---|---|---|---|---|---|---|---|---|---|---|---|
| | WT | SctG-sfGFP | | | WT 1 | WT 2 | WT 3 | SctG-sfGFP 1 | SctG-sfGFP 2 | SctG-sfGFP 3 | |
| Nuclease SbcCD subunit D | 6.08E+06 | 8.57E+07 | 3.821 | 0.0000 | 6.2E+06 | 6.6E+06 | 5.4E+06 | 8.6E+07 | 8.6E+07 | 8.5E+07 | 3 |
| Putative uncharacterized protein | 5.96E+06 | 4.40E+07 | 2.903 | 0.0000 | 5.2E+06 | 7.5E+06 | 5.3E+06 | 4.2E+07 | 4.3E+07 | 4.7E+07 | 3 |
| Phosphoenolpyruvate-protein phosphotransferase PTSP | 1.34E+07 | 3.31E+07 | 1.371 | 0.0094 | 1.1E+07 | 2.0E+07 | 9.0E+06 | 3.4E+07 | 2.7E+07 | 3.9E+07 | 9 |
| T3SS component SctD | 8.28E+07 | 1.66E+08 | 1.012 | 0.0001 | 9.2E+07 | 7.3E+07 | 8.3E+07 | 1.7E+08 | 1.6E+08 | 1.7E+08 | 13 |
| T3SS effector YopH | 6.08E+08 | 1.18E+09 | 0.957 | 0.0000 | 6.1E+08 | 6.1E+08 | 6.0E+08 | 1.1E+09 | 1.3E+09 | 1.1E+09 | 32 |
| T3SS lipoprotein component SctG (YscW) | 2.24E+08 | 4.28E+08 | 0.934 | 0.0000 | 2.1E+08 | 2.3E+08 | 2.3E+08 | 4.2E+08 | 4.3E+08 | 4.4E+08 | 9 |
| T3SS component SctC | 1.61E+08 | 2.80E+08 | 0.793 | 0.0000 | 1.6E+08 | 1.7E+08 | 1.6E+08 | 2.6E+08 | 2.9E+08 | 2.8E+08 | 20 |
| T3SS chaperone YscG | 4.23E+08 | 6.85E+08 | 0.700 | 0.0007 | 4.4E+08 | 4.5E+08 | 3.7E+08 | 6.6E+08 | 6.8E+08 | 7.2E+08 | 5 |
| T3SS chaperone YscE | 4.23E+08 | 6.70E+08 | 0.666 | 0.0001 | 4.4E+08 | 4.4E+08 | 3.9E+08 | 6.5E+08 | 6.7E+08 | 6.9E+08 | 7 |
| T3SS component SctF | 3.76E+08 | 5.94E+08 | 0.658 | 0.0000 | 3.7E+08 | 3.9E+08 | 3.6E+08 | 5.9E+08 | 6.0E+08 | 5.9E+08 | 4 |
| T3SS gatekeeper SctW (YopN) | 2.24E+08 | 3.52E+08 | 0.654 | 0.0001 | 2.1E+08 | 2.4E+08 | 2.2E+08 | 3.6E+08 | 3.4E+08 | 3.6E+08 | 5 |
| T3SS component SctJ | 2.72E+08 | 4.13E+08 | 0.606 | 0.0001 | 2.7E+08 | 2.9E+08 | 2.6E+08 | 4.1E+08 | 4.2E+08 | 4.1E+08 | 13 |
| T3SS regulator YscM1 | 4.28E+08 | 6.33E+08 | 0.568 | 0.0008 | 4.4E+08 | 4.6E+08 | 3.8E+08 | 6.3E+08 | 6.4E+08 | 6.3E+08 | 8 |
| T3SS protein YscB | 4.84E+08 | 6.85E+08 | 0.506 | 0.0011 | 4.4E+08 | 5.3E+08 | 4.8E+08 | 6.8E+08 | 6.9E+08 | 6.8E+08 | 4 |
| T3SS component YscX | 6.45E+07 | 8.92E+07 | 0.468 | 0.0004 | 6.5E+07 | 6.6E+07 | 6.2E+07 | 9.3E+07 | 9.1E+07 | 8.4E+07 | 3 |
| T3SS component SctL | 3.28E+08 | 4.55E+08 | 0.466 | 0.0032 | 3.1E+08 | 3.3E+08 | 3.5E+08 | 4.1E+08 | 4.9E+08 | 4.7E+08 | 9 |
| T3SS effector YopM | 3.86E+08 | 5.13E+08 | 0.410 | 0.0002 | 3.8E+08 | 3.9E+08 | 3.8E+08 | 5.1E+08 | 5.0E+08 | 5.3E+08 | 3 |
| Protein translocase subunit SecA | 3.78E+08 | 4.96E+08 | 0.391 | 0.0003 | 3.8E+08 | 3.8E+08 | 3.7E+08 | 4.8E+08 | 5.1E+08 | 4.9E+08 | 44 |
| T3SS component SctN | 1.06E+08 | 1.36E+08 | 0.363 | 0.0016 | 1.1E+08 | 1.1E+08 | 1.0E+08 | 1.3E+08 | 1.4E+08 | 1.4E+08 | 17 |
| T3SS chaperone SycD | 1.66E+08 | 2.10E+08 | 0.342 | 0.0054 | 1.8E+08 | 1.6E+08 | 1.6E+08 | 2.0E+08 | 2.1E+08 | 2.2E+08 | 6 |

| | | | | | | | | | | | |
|---|---|---|---|---|---|---|---|---|---|---|---|
| T3SS chaperone SycN | 4.43E+08 | 5.41E+08 | 0.288 | 0.0058 | 4.6E+08 | 4.6E+08 | 4.1E+08 | 5.4E+08 | 5.3E+08 | 5.5E+08 | 5 |
| SsrA-binding protein | 8.01E+07 | 9.71E+07 | 0.277 | 0.0072 | 8.0E+07 | 8.2E+07 | 7.9E+07 | 9.9E+07 | 1.0E+08 | 9.0E+07 | 3 |
| T3SS translocator SctB (YopD) | 1.11E+09 | 1.33E+09 | 0.259 | 0.0074 | 1.0E+09 | 1.2E+09 | 1.1E+09 | 1.3E+09 | 1.4E+09 | 1.3E+09 | 14 |
| Putative NADP-dependent oxidoreductase yncB | 1.80E+08 | 1.56E+08 | -0.209 | 0.0074 | 1.7E+08 | 1.9E+08 | 1.8E+08 | 1.6E+08 | 1.5E+08 | 1.6E+08 | 17 |
| Catalase | 4.63E+08 | 3.95E+08 | -0.228 | 0.0076 | 4.4E+08 | 4.7E+08 | 4.8E+08 | 3.9E+08 | 3.9E+08 | 4.1E+08 | 18 |
| Urease accessory protein UreD | 1.59E+08 | 1.29E+08 | -0.303 | 0.0020 | 1.6E+08 | 1.7E+08 | 1.5E+08 | 1.3E+08 | 1.3E+08 | 1.3E+08 | 10 |
| 3,4-dihydroxy-2-butanone 4-phosphate synthase | 1.56E+08 | 1.26E+08 | -0.307 | 0.0091 | 1.4E+08 | 1.7E+08 | 1.6E+08 | 1.2E+08 | 1.3E+08 | 1.2E+08 | 7 |
| Putative uncharacterized protein | 1.16E+08 | 9.26E+07 | -0.325 | 0.0032 | 1.2E+08 | 1.2E+08 | 1.1E+08 | 9.0E+07 | 9.6E+07 | 9.2E+07 | 5 |
| Uncharacterized protein yibQ | 2.30E+07 | 1.77E+07 | -0.378 | 0.0090 | 2.3E+07 | 2.4E+07 | 2.2E+07 | 1.6E+07 | 1.9E+07 | 1.8E+07 | 3 |
| Putative uncharacterized protein | 8.35E+07 | 6.28E+07 | -0.410 | 0.0036 | 8.1E+07 | 7.8E+07 | 9.1E+07 | 6.7E+07 | 6.1E+07 | 6.1E+07 | 5 |
| Nitrate/nitrite response regulator protein narP | 8.58E+07 | 5.63E+07 | -0.606 | 0.0014 | 7.8E+07 | 8.4E+07 | 9.5E+07 | 5.6E+07 | 6.0E+07 | 5.3E+07 | 7 |
| Putative uncharacterized protein | 1.22E+08 | 7.01E+07 | -0.808 | 0.0015 | 1.2E+08 | 1.2E+08 | 1.2E+08 | 8.4E+07 | 6.0E+07 | 6.7E+07 | 3 |
| ATP-dependent DNA helicase recQ | 1.29E+07 | 6.94E+06 | -0.892 | 0.0005 | 1.2E+07 | 1.2E+07 | 1.5E+07 | 6.6E+06 | 7.6E+06 | 6.6E+06 | 5 |
| Cytochrome c-type biogenesis protein CcmE | 1.72E+07 | 5.45E+06 | -1.652 | 0.0037 | 1.1E+07 | 2.0E+07 | 2.0E+07 | 4.6E+06 | 4.4E+06 | 7.4E+06 | 5 |
| Molybdopterin biosynthesis protein moeB | 2.02E+07 | 5.59E+06 | -1.731 | 0.0080 | 2.6E+07 | 2.5E+07 | 9.8E+06 | 5.0E+06 | 6.4E+06 | 5.4E+06 | 4 |
| Selenocysteine-specific elongation factor | 7.39E+07 | 7.17E+06 | -3.068 | 0.0097 | 9.6E+07 | 1.9E+07 | 1.1E+08 | 7.1E+06 | 4.7E+06 | 9.7E+06 | 4 |
| T3SS transcriptional activator VirF | 4.58E+07 | 6.99E+07 | 0.629 | 0.0122 | 5.1E+07 | 5.1E+07 | 3.5E+07 | 6.5E+07 | 7.1E+07 | 7.4E+07 | 4 |

**Table S5 – Specific interactors of SctG**

Proteins detected by quantitative mass spectrometry after co-immunoprecipitation with anti-GFP beads in *Y. enterocolitica* strains expressing SctG-sfGFP from its native location on the virulence plasmid (SctG-sfGFP) and a control expressing cytosolic sfGFP from plasmid at a comparable level (sfGFP).

All proteins with $p < 0.001$ and number of detected peptides (# pept.) > 2 are included in the list, and ordered according to their enrichment in the SctG-sfGFP co-immunoprecipitation sample. SctG is shaded in orange, other T3SS components are shaded in yellow. Significant interactors discussed in the main text are shaded in blue.

| Protein | Mean intensity | | log 2 intensity ratio | $-\log_{10}$ ($p$ val.) | Individual intensities | | | | | | # pept. |
|---|---|---|---|---|---|---|---|---|---|---|---|
| | SctG-sfGFP | sfGFP | | | SctG-sfGFP 1 | SctG-sfGFP 2 | SctG-sfGFP 3 | sfGFP 1 | sfGFP 2 | sfGFP 3 | |
| PTS trehalose transporter subunit EIIBC | 2.41E+09 | 5.94E+05 | 11.985 | 6.71 | 2.4E+09 | 1.5E+09 | 3.3E+09 | 3.8E+05 | 5.5E+05 | 8.5E+05 | 9 |
| T3SS lipoprotein component SctG (YscW) | 7.14E+09 | 1.43E+07 | 8.956 | 7.36 | 7.0E+09 | 5.6E+09 | 8.8E+09 | 1.1E+07 | 1.6E+07 | 1.6E+07 | 11 |
| T3SS component SctC | 7.07E+08 | 1.26E+07 | 5.806 | 6.70 | 7.5E+08 | 5.7E+08 | 8.0E+08 | 1.1E+07 | 1.5E+07 | 1.2E+07 | 35 |
| T3SS effector YopM | 3.59E+08 | 1.19E+07 | 4.878 | 5.22 | 5.0E+08 | 2.5E+08 | 3.3E+08 | 1.2E+07 | 1.4E+07 | 9.2E+06 | 4 |
| PTS beta-glucoside transporter subunit IIABC | 2.24E+07 | 8.06E+05 | 4.874 | 4.34 | 1.9E+07 | 1.7E+07 | 3.1E+07 | 4.0E+05 | 1.1E+06 | 9.4E+05 | 5 |
| Lon | 3.65E+07 | 1.86E+06 | 4.373 | 4.05 | 4.2E+07 | 2.4E+07 | 4.3E+07 | 2.5E+06 | 2.2E+06 | 9.2E+05 | 18 |
| Rib. prot. L11 | 6.00E+07 | 3.66E+06 | 4.059 | 4.82 | 5.3E+07 | 5.0E+07 | 7.8E+07 | 4.9E+06 | 3.5E+06 | 2.5E+06 | 13 |
| K+ channel | 1.19E+08 | 9.45E+06 | 3.629 | 4.65 | 9.9E+07 | 9.3E+07 | 1.6E+08 | 1.2E+07 | 7.9E+06 | 8.4E+06 | 8 |
| hypothetical protein CBX71048 | 4.74E+06 | 6.51E+05 | 3.058 | 3.24 | 5.0E+06 | 4.4E+06 | 4.8E+06 | 1.1E+06 | 3.5E+05 | 4.5E+05 | 7 |
| T3SS component SctV | 2.29E+08 | 3.73E+07 | 2.630 | 5.07 | 2.5E+08 | 2.0E+08 | 2.4E+08 | 3.1E+07 | 4.3E+07 | 3.8E+07 | 23 |
| LolE | 6.42E+06 | 1.14E+06 | 2.452 | 3.18 | 5.5E+06 | 4.3E+06 | 9.5E+06 | 1.1E+06 | 8.1E+05 | 1.5E+06 | 5 |
| FtsZ | 3.19E+08 | 6.65E+07 | 2.256 | 4.92 | 2.9E+08 | 2.9E+08 | 3.8E+08 | 5.9E+07 | 6.9E+07 | 7.1E+07 | 31 |
| Coproporphyrinogen-III oxidase hemN | 7.07E+06 | 1.55E+06 | 2.245 | 3.82 | 7.4E+06 | 6.8E+06 | 7.0E+06 | 1.3E+06 | 1.2E+06 | 2.2E+06 | 16 |
| RNA polymerase sigma-54 factor rpoN | 5.36E+06 | 1.12E+06 | 2.228 | 3.31 | 7.3E+06 | 5.1E+06 | 3.7E+06 | 1.4E+06 | 1.1E+06 | 8.5E+05 | 8 |
| Hypothetical protein CBX71617 | 7.26E+06 | 1.53E+06 | 2.207 | 3.59 | 7.2E+06 | 5.0E+06 | 9.6E+06 | 1.5E+06 | 1.3E+06 | 1.8E+06 | 3 |
| Uncharacterized protein yjaG | 1.03E+08 | 2.31E+07 | 2.143 | 4.00 | 8.6E+07 | 9.1E+07 | 1.3E+08 | 1.9E+07 | 2.3E+07 | 2.7E+07 | 10 |
| Hypothetical protein CBX72046 | 3.51E+07 | 8.00E+06 | 2.117 | 3.38 | 2.6E+07 | 3.2E+07 | 4.7E+07 | 9.8E+06 | 6.0E+06 | 8.2E+06 | 14 |
| Inner membrane protein ytfL | 1.98E+07 | 5.44E+06 | 1.863 | 4.42 | 2.2E+07 | 1.7E+07 | 1.9E+07 | 4.7E+06 | 6.3E+06 | 5.3E+06 | 9 |
| Ubiquinol oxidase subunit 1 cyoB | 1.00E+08 | 2.81E+07 | 1.806 | 3.09 | 9.6E+07 | 7.0E+07 | 1.3E+08 | 2.5E+07 | 3.5E+07 | 2.5E+07 | 7 |

| | | | | | | | | | | | |
|---|---|---|---|---|---|---|---|---|---|---|---|
| Hydrogenase isoenzymes nickel incorporation protein hypB | 2.11E+07 | 6.73E+06 | 1.679 | 3.20 | 1.9E+07 | 1.9E+07 | 2.5E+07 | 5.4E+06 | 5.7E+06 | 9.1E+06 | 9 |
| Protein-export membrane protein SecF | 6.34E+07 | 2.35E+07 | 1.454 | 3.35 | 6.7E+07 | 5.7E+07 | 6.6E+07 | 1.8E+07 | 2.4E+07 | 2.8E+07 | 8 |
| Proline/betaine transporter proP | 2.82E+07 | 1.07E+07 | 1.387 | 3.75 | 3.0E+07 | 2.4E+07 | 3.1E+07 | 9.5E+06 | 1.2E+07 | 1.1E+07 | 8 |
| Glycine cleavage system transcriptional repressor gcvR | 2.50E+06 | 1.07E+06 | 1.235 | 3.29 | 2.8E+06 | 2.5E+06 | 2.2E+06 | 1.1E+06 | 1.2E+06 | 8.7E+05 | 3 |
| ATP synthase subunit alpha | 2.78E+08 | 1.27E+08 | 1.128 | 3.79 | 3.0E+08 | 2.6E+08 | 2.8E+08 | 1.4E+08 | 1.2E+08 | 1.2E+08 | 30 |
| Anaerobic ribonucleoside-triphosphate reductase nrdD | 3.79E+07 | 1.85E+07 | 1.045 | 3.18 | 3.9E+07 | 3.8E+07 | 3.6E+07 | 2.1E+07 | 1.5E+07 | 2.0E+07 | 21 |
| ATP synthase subunit beta | 2.63E+08 | 1.32E+08 | 0.993 | 3.69 | 2.8E+08 | 2.5E+08 | 2.6E+08 | 1.4E+08 | 1.3E+08 | 1.2E+08 | 26 |
| Uncharacterized protein ytfN | 7.31E+06 | 3.72E+06 | 0.982 | 3.13 | 8.0E+06 | 7.0E+06 | 7.0E+06 | 3.1E+06 | 3.9E+06 | 4.1E+06 | 31 |
| Alpha-1,4 glucan phosphorylase | 2.08E+07 | 1.07E+07 | 0.960 | 3.12 | 1.8E+07 | 2.0E+07 | 2.4E+07 | 1.0E+07 | 1.2E+07 | 9.9E+06 | 4 |
| ATP synthase gamma chain | 4.91E+07 | 2.56E+07 | 0.942 | 3.40 | 5.4E+07 | 4.6E+07 | 4.8E+07 | 2.8E+07 | 2.4E+07 | 2.4E+07 | 22 |
| Lipoprotein YlpA | 8.53E+07 | 4.69E+07 | 0.866 | 3.06 | 9.3E+07 | 8.4E+07 | 7.9E+07 | 5.3E+07 | 4.6E+07 | 4.2E+07 | 6 |
| Succinate dehydrogenase iron-sulfur subunit sdhB | 6.65E+06 | 3.69E+06 | 0.852 | 3.00 | 5.9E+06 | 6.9E+06 | 7.1E+06 | 3.6E+06 | 4.1E+06 | 3.3E+06 | 4 |
| Adhesin YadA | 6.01E+08 | 3.37E+08 | 0.836 | 3.06 | 6.3E+08 | 6.3E+08 | 5.4E+08 | 3.2E+08 | 3.7E+08 | 3.2E+08 | 24 |
| Protein translocase subunit SecA | 4.66E+07 | 2.96E+07 | 0.655 | 3.15 | 4.6E+07 | 4.5E+07 | 4.9E+07 | 3.0E+07 | 3.0E+07 | 2.9E+07 | 43 |
| 50S ribosomal protein L4 rplD | 6.70E+08 | 4.29E+08 | 0.644 | 3.15 | 6.6E+08 | 6.5E+08 | 6.9E+08 | 4.3E+08 | 4.3E+08 | 4.3E+08 | 11 |
| Pyridoxine 5-phosphate synthase pdxJ | 1.55E+07 | 2.64E+07 | -0.765 | 3.27 | 1.5E+07 | 1.5E+07 | 1.7E+07 | 2.5E+07 | 2.8E+07 | 2.7E+07 | 3 |
| Uncharacterized protein yigL | 1.87E+06 | 3.23E+06 | -0.788 | 3.06 | 1.8E+06 | 2.0E+06 | 1.8E+06 | 3.5E+06 | 3.3E+06 | 2.9E+06 | 6 |
| 3-mercaptopyruvate sulfurtransferase sseA | 4.67E+06 | 8.44E+06 | -0.854 | 3.07 | 4.9E+06 | 4.2E+06 | 5.0E+06 | 7.6E+06 | 9.0E+06 | 8.8E+06 | 4 |
| Methionine aminopeptidase map | 9.85E+06 | 1.90E+07 | -0.941 | 3.36 | 9.4E+06 | 1.1E+07 | 9.6E+06 | 2.1E+07 | 1.9E+07 | 1.7E+07 | 13 |
| Glycine betaine-binding periplasmic protein proX | 5.82E+06 | 1.15E+07 | -0.986 | 3.98 | 5.9E+06 | 5.7E+06 | 5.9E+06 | 1.1E+07 | 1.2E+07 | 1.1E+07 | 12 |
| Outer membrane permeability protein sanA | 6.31E+05 | 1.31E+06 | -1.054 | 3.11 | 6.6E+05 | 6.9E+05 | 5.4E+05 | 1.3E+06 | 1.1E+06 | 1.5E+06 | 3 |
| 30S ribosomal protein S19 | 1.46E+08 | 3.17E+08 | -1.130 | 3.08 | 1.6E+08 | 1.1E+08 | 1.6E+08 | 3.4E+08 | 3.0E+08 | 3.1E+08 | 5 |
| YopH effector chaperone SycH | 4.12E+07 | 9.13E+07 | -1.144 | 4.12 | 4.1E+07 | 4.2E+07 | 4.0E+07 | 8.2E+07 | 9.6E+07 | 9.5E+07 | 10 |
| Hypothetical protein CBX70265 | 2.39E+06 | 5.33E+06 | -1.157 | 3.60 | 2.2E+06 | 2.3E+06 | 2.7E+06 | 6.0E+06 | 5.2E+06 | 4.8E+06 | 3 |
| UPF0227 protein CBX70961 | 3.36E+06 | 7.72E+06 | -1.199 | 3.60 | 3.3E+06 | 3.8E+06 | 3.0E+06 | 8.2E+06 | 6.7E+06 | 8.3E+06 | 9 |
| Protein ygiW | 1.48E+07 | 3.75E+07 | -1.349 | 3.61 | 1.2E+07 | 1.5E+07 | 1.7E+07 | 4.0E+07 | 3.3E+07 | 3.9E+07 | 6 |

| | | | | | | | | | | | |
|---|---|---|---|---|---|---|---|---|---|---|---|
| Formate--tetrahydrofolate ligase fhs | 8.75E+05 | 2.48E+06 | -1.524 | 3.57 | 7.1E+05 | 1.1E+06 | 8.3E+05 | 2.2E+06 | 2.6E+06 | 2.6E+06 | 4 |
| Hypothetical protein CBX71612 | 2.48E+06 | 1.06E+07 | -2.096 | 5.73 | 2.4E+06 | 2.6E+06 | 2.5E+06 | 1.1E+07 | 1.0E+07 | 1.0E+07 | 5 |
| Putative uncharacterized protein ybcY | 6.67E+05 | 2.84E+06 | -2.118 | 3.55 | 5.9E+05 | 5.0E+05 | 9.1E+05 | 2.4E+06 | 3.5E+06 | 2.6E+06 | 4 |
| Ribonucleoside-diphosphate reductase | 1.06E+06 | 5.14E+06 | -2.290 | 4.12 | 8.7E+05 | 1.4E+06 | 9.5E+05 | 6.2E+06 | 4.9E+06 | 4.3E+06 | 8 |
| L-ribulose-5-phosphate 4-epimerase araD | 7.71E+05 | 3.87E+06 | -2.365 | 3.09 | 4.7E+05 | 7.5E+05 | 1.1E+06 | 3.3E+06 | 5.2E+06 | 3.0E+06 | 4 |
| Protein ychN | 1.01E+06 | 5.51E+06 | -2.548 | 3.23 | 1.0E+06 | 5.5E+05 | 1.4E+06 | 5.7E+06 | 6.3E+06 | 4.6E+06 | 3 |
| Chelated iron transport system membrane protein yfeB | 1.13E+06 | 8.32E+06 | -2.972 | 3.04 | 1.4E+06 | 1.5E+06 | 5.2E+05 | 6.2E+06 | 1.1E+07 | 7.3E+06 | 7 |
| Hypothetical protein CBX70345 | 1.11E+06 | 9.51E+06 | -3.226 | 3.58 | 1.8E+06 | 6.8E+05 | 8.1E+05 | 7.8E+06 | 9.3E+06 | 1.2E+07 | 12 |
| Hypothetical protein CBX73386 | 1.02E+06 | 2.33E+07 | -4.588 | 4.56 | 1.6E+06 | 6.6E+05 | 8.2E+05 | 1.7E+07 | 2.4E+07 | 2.9E+07 | 3 |
| Hypothetical protein CBX71757 | 2.00E+06 | 6.93E+07 | -5.128 | 6.96 | 2.4E+06 | 1.9E+06 | 1.7E+06 | 6.3E+07 | 7.3E+07 | 7.1E+07 | 4 |
| Hypothetical protein CBX74219 | 1.47E+06 | 5.31E+07 | -5.299 | 4.91 | 1.3E+06 | 2.3E+06 | 8.3E+05 | 5.0E+07 | 5.2E+07 | 5.8E+07 | 3 |
| Hypothetical protein CBX73387 | 6.74E+05 | 4.75E+07 | -6.075 | 4.64 | 5.9E+05 | 9.8E+05 | 4.6E+05 | 2.3E+07 | 5.3E+07 | 6.7E+07 | 9 |
| UDP-3-O-(3-hydroxymyristoyl) glucosamine N-acyltransferase lpxD | 5.18E+06 | 8.35E+08 | -7.210 | 5.61 | 4.6E+06 | 5.3E+06 | 5.7E+06 | 4.4E+08 | 8.0E+08 | 1.3E+09 | 14 |

**Table S6 - PTS trehalose transporter subunit EIIBC has an influence on the expression of OmpW and SctG, but very few other proteins**

Comparison of expression levels in total cellular protein samples of wild-type (WT) and *ΔptsEIIBC* (dPTS) *Y. enterocolitica* under non-secreting and secreting conditions (ns, s), determined by label-free mass spectrometry. Light grey font, imputed values (no peptides detected in at least one sample).

All proteins with an average log 2 intensity ratio (WT / *ΔptsEIIBC*) of > 1 and number of detected peptides (# pept.) > 2 are included in the list, and ordered according to their intensity ratio WT / *ΔptsEIIBC*. SctG is shaded in orange, OmpW in green.

| **Protein** | **avg. log 2 intensity ratio** | **Individual intensities** | | | | **# pept.** |
|---|---|---|---|---|---|---|
| | | **WT ns** | **dPTS ns** | **WT secr** | **dPTS secr** | |
| PTS trehalose transporter subunit EIIBC | 2.54 | 2.45E+07 | 3.92E+06 | 2.80E+07 | 5.17E+06 | 3 |
| Ornithine decarboxylase, inducible | 3.52 | 2.21E+07 | 3.98E+06 | 4.31E+07 | 1.81E+06 | 12 |
| Outer membrane protein W | 2.61 | 4.18E+08 | 7.02E+07 | 4.78E+08 | 7.66E+07 | 6 |
| Ribosomal-protein-alanine acetyltransferase | 2.30 | 3.23E+07 | 5.23E+06 | 1.03E+07 | 2.63E+06 | 3 |
| Anaerobic ribonucleoside-triphosphate reductase | 1.81 | 2.26E+08 | 6.73E+07 | 2.29E+08 | 6.25E+07 | 11 |
| Hydrogenase-2 large chain | 1.54 | 4.27E+07 | 1.51E+07 | 4.87E+07 | 1.63E+07 | 13 |
| L-asparaginase 2 | 1.52 | 2.72E+08 | 1.03E+08 | 3.11E+08 | 9.98E+07 | 10 |
| Hydrogenase isoenzymes nickel incorporation protein hypB | 1.35 | 2.85E+07 | 1.12E+07 | 2.93E+07 | 1.15E+07 | 3 |
| T3SS lipoprotein component SctG (YscW) | 1.29 | 2.22E+08 | 9.28E+07 | 2.37E+08 | 9.50E+07 | 7 |
| 5-methyltetrahydropteroyltriglutamate-homocysteine methyltransferase | 1.20 | 2.44E+08 | 1.12E+08 | 2.83E+08 | 1.16E+08 | 28 |
| Putative phosphotransferase YE2177 (Fragment) | 1.15 | 7.68E+06 | 6.36E+06 | 1.50E+07 | 3.67E+06 | 3 |

**Table S7 – Strains and plasmids used in this study**

Except for pSW048, all plasmids encoding T3SS components use the *Yersinia enterocolitica* variant.

| Strain | Genotype | Reference |
|---|---|---|
| MRS40 | Wild-type *Y. enterocolitica* E40 *ΔblaA* | [7] |
| IML421*asd* (ΔHOPEMTasd) | MRS40 $yopO_{\Delta 12\text{-}427}$ $yopE_{21}$ $yopH_{\Delta 11\text{-}352}$ $yopM_{23}$ $yopP_{23}$ $yopT_{135}$ *Δasd* | [8] |
| pYV⁻ | MRS40 cured of its natural virulence plasmid | |
| AD4016 | MRS40 *egfp-sctQ* | [9] |
| AD4051 | MRS40 Δ*sctD* | [9] |
| AD4085 | IML421*asd egfp-sctQ* | [8] |
| AD4175 | IML421*asd sctV-egfp* | [10] |
| AD4306 | IML421*asd egfp-sctD* | [11] |
| AD4393 | IML421*asd mCherry-sctD egfp-sctQ* | [12] |
| AD4411 | IML421*asd egfp-sctQ ΔsctD* | [12] |
| ADMA4098 | MRS40 Δ*sctC* | This work |
| ADTM4520 | IML421*asd egfp-sctL* | [12] |
| ADTM4521 | IML421*asd mCherry-sctL* | [12] |
| CB4001 | MRS40 *sctC-mCherry* Δ*sctG* | This work |
| CH4008 | MRS40 *sctC-mCherry* sctG-*sfgfp* | This work |
| CH4010 | MRS40 *mCherry-sctL* sctG-*sfgfp* | This work |
| CHSW4001 | MRS40 *mCherry-sctL* | This work |
| MA4005 | MRS40 *sctC-mCherry* | [9] |
| MF4002 | MRS40 *sctG-sfgfp* | This work |
| MF4003 | MRS40 *mCherry-SctD* sctG-*sfgfp* | This work |
| MF4005 | MRS40 Δ*sctG* | This work |
| MF015 | MRS40 Δ*sctG mCherry-sctL* | This work |
| MF4018 | IML421*asd* sctG-*sfgfp* | This work |
| MF4015 | MRS40 *egfp-sctL* Δ*sctG* | This work |
| SW4062 | MRS40 Δ*pts* | This work |
| SW4064 | MRS40 Δ*pts* Δ*sctG* | This work |
| SW4080 | MRS40 *sctV-egfp* Δ*sctG* | This work |
| SW4081 | MRS40 *sctV-egfp* Δ*sctC* | This work |

| Plasmids | Genotype | Reference |
|---|---|---|
| pBAD-His B | pBR322-derived expression vector | Invitrogen |
| pKNG101 | *oriR6K sacBR⁺ oriTRK2 strAB⁺* (suicide vector for homologous recombination) | [13] |
| pAD208 | *pKNG101-sctV-egfp* | [12] |
| pAD309 | *pKNG-mCherry-sctD* | This work |
| pAD476 | *pBAD::egfp* | This work |
| pAD638 | $pBAD::sctF_{S5C}$ | [10] |
| pADTM511 | *pKNG101-egfp-sctL* | [12] |
| pCH03 | *pKNG101-sctG-sfgfp* | This work |
| pCL5 | *oripBR322, oriTRK2,* $p_{tac}$*::virF* | [14] |
| pMA12 | p*KNG101-sctC-mCherry* | [9] |
| pMA26 | p*KNG101-ΔsctC* | [9] |

| | | |
|---|---|---|
| pMF001 | *pBAD::sctG* | This work |
| pMF002 | *pBAD::sctG-sfgfp* | This work |
| pMF003 | *pKNG101-ΔsctG* | This work |
| pMF012 | *pKNG101-ΔptsIIbc* | This work |
| pSW048 | *pBAD::Pa_sctG (P. aeruginosa exsB)* | This work |
| pSW066 | *pBAD::ptsIIbc-flag* | This work |

**Text S1 – Extended discussion of interaction studies**

The strongest interaction with the pilotin protein found in our study was the highly specific enrichment of several PTS sugar transporter subunits, most significantly the membrane-bound trehalose transporter subunit EIIBC (Fig. 3, Table S5). Indeed, PTS systems were repeatedly associated with bacterial virulence (reviewed in [15,16]): amongst others, presence of the PTS glucose transporter subunit EIIBC conferred a competitive advantage to *Y. pestis* in serum-like medium [17], a *Vibrio cholerae* PTS system was found to modulate virulence gene expression [18], and *Salmonella* Typhimurium lacking a PTS EI subunit showed a nearly 1000-fold increase in $LD_{50}$ [19]. Specifically a PTS trehalose transporter system induces hypervirulence in *Clostridium difficile* [20]. Notably, the PTS glucose transporter EIIA subunit directly interacts with cytosolic components of the *Salmonella* SPI-2 T3SS and activates type III secretion [21]. Our experiments failed to show a direct link between PTS II BC and T3SS assembly and activity. Overall, the two most likely links between PTS sugar transporters and virulence appear to be energy provision and/or host sensing; further experiments are required to describe the connection in more detail.

Notably, while the pilotin mostly fractionates with the OM in membrane preparations [22–24], many of the proteins found to interact with the labeled pilotin in the co-immunoprecipitation assay are integral IM proteins or IM-associated proteins. The used SctG-sfGFP fusion is expressed from its native genetic location, supports native secretion (Fig. S4), and interacts with both known interactors, the secretin SctC and proteins of the Lol pathway, in our assays (Fig. 3). While these are strong indications for correct localization and interactions of SctG-sfGFP, it is possible that transport of the fusion protein to the OM is impaired (or the observed slightly increased levels of SctG-sfGFP saturate the export pathways to the OM), which in turn may enhance the fraction of pilotins that can contact these IM(-associated) proteins. Alternatively, the interactions may be caused by natively soluble pilotin proteins, possibly bound to other proteins such as LolB [25,26]. Finally, the N-terminal lipid anchor of the pilotin might be removed after assembly of the secretin ring or the injectisome, which would render the protein soluble [23] in the periplasm.

Other significant interaction partners of the pilotin outside the T3SS are FtsZ, a prokaryotic tubulin homolog with a key role in cell division, the protease Lon, components of the ATP synthase (subunits $\alpha$, $\beta$, $\gamma$), the Sec export pathway (SecF/A), and LolD/E, two components of the ABC transporter of the Lol lipoprotein export pathway in the IM. FtsZ is an interesting interactor, as it might play a role in the long-known, but still incompletely understood link between T3SS activity and cell growth and division (actively secreting bacteria of many species cease growth and division and are often bigger than T3SS-deficient or non-secreting bacteria [27–30]). Since to our knowledge, this is the first time that a T3SS component was found to be associated with bacterial cytoskeleton proteins, this might indicate an involvement of SctG in the non-random localization of T3SS around the cell [31,32]. The Lon protease degrades unfolded or misfolded proteins, especially under stress conditions [33], but can also have more specific roles in metabolism, cell cycle control and virulence through proteolysis of specific regulatory proteins [34,35]. While it cannot be ruled out that the specific interaction of SctG-sfGFP with Lon denotes degradation of partially misfolded protein, Lon proteases have been shown to influence T3SS activity, both positively via degradation of histone-like proteins like *Yersinia* YmoA that suppress the transcription of T3SS components [36–39] and negatively, most likely by directly degrading key components of the T3SS, mainly in plant pathogens [40–43].

**<u>Description of additional supporting files</u>**

**Movie S1**

Time-lapse microscopy video of live *Y. enterocolitica* SctG-sfGFP mCherry-SctD on an agarose pad. Green and red fluorescence channel as well as an overlay of both channels are displayed (red channel displayed in magenta for the overlay). Images were taken every 2 s for a total of 58 s as indicated. Scale bar, 2 µm.

**Dataset S1: The pilotin protein influences assembly of all tested injectisome components**

Large fields of view of localization of mCherry-SctL and $SctF_{S5C}$, stained with maleimide CF 488A, in wild-type background and absence of SctG. For $SctF_{S5C}$, a deletion of *sctD* was used as a negative control and foci counts were corrected for this negative control. Fractions of bacteria displaying foci in the needle staining are indicated in the dataset for each individual field of view (three fields of view (fov) each for 3-6 independent experiments).

**References for Supporting Information**

1. Wallace IM, O'Sullivan O, Higgins DG, Notredame C. M-Coffee: combining multiple sequence alignment methods with T-Coffee. Nucleic Acids Res. 2006;34:1692–9.
2. Walker KA, Miller VL. Synchronous Gene Expression of the Yersinia enterocolitica Ysa Type III Secretion System and Its Effectors. J Bacteriol. 2009 Mar 15;191(6):1816–26.
3. Hueck CJ. Type III protein secretion systems in bacterial pathogens of animals and plants. Microbiol Mol Biol Rev. 1998 Jun;62(2):379–433.
4. Portaliou AG, Tsolis KC, Loos MS, Zorzini V, Economou A. Type III Secretion: Building and Operating a Remarkable Nanomachine. Trends Biochem Sci. 2016 Feb;41(2):175–89.
5. Wagner S, Diepold A. A Unified Nomenclature for Injectisome-Type Type III Secretion Systems. In: Current topics in microbiology and immunology. 2020. p. 427: 1-10.
6. Diepold A, Wagner S. Assembly of the bacterial type III secretion machinery. FEMS Microbiol Rev. 2014 Jul 3;38(4):802–22.
7. Sory M-P, Boland A, Lambermont I, Cornelis GR. Identification of the YopE and YopH domains required for secretion and internalization into the cytosol of macrophages, using the cyaA gene fusion approach. Proc Natl Acad Sci U S A. 1995 Dec 18;92(26):11998–2002.
8. Kudryashev M, Stenta M, Schmelz S, Amstutz M, Wiesand U, Castaño-Díez D, et al. In situ structural analysis of the Yersinia enterocolitica injectisome. Elife. 2013 Jul 30;2(Im):e00792.
9. Diepold A, Amstutz M, Abel S, Sorg I, Jenal U, Cornelis GR. Deciphering the assembly of the Yersinia type III secretion injectisome. EMBO J. 2010 Jun 2;29(11):1928–40.
10. Wimmi S, Balinovic A, Jeckel H, Selinger L, Lampaki D, Eisemann E, et al. Dynamic relocalization of cytosolic type III secretion system components prevents premature protein secretion at low external pH. Nat Commun. 2021 Dec 12;12(1):1625.
11. Diepold A, Kudryashev M, Delalez NJ, Berry RM, Armitage JP. Composition, Formation, and Regulation of the Cytosolic C-ring, a Dynamic Component of the Type III Secretion Injectisome. Stock AM, editor. PLOS Biol. 2015 Jan 15;13(1):e1002039.
12. Diepold A, Sezgin E, Huseyin M, Mortimer T, Eggeling C, Armitage JP. A dynamic and adaptive network of cytosolic interactions governs protein export by the T3SS injectisome. Nat Commun. 2017 Aug 27;8(1):15940.
13. Kaniga K, Delor I, Cornelis GR. A wide-host-range suicide vector for improving reverse genetics in Gram-negative bacteria: inactivation of the blaA gene of Yersinia enterocolitica. Gene. 1991 Dec 20;109(1):137–41.
14. Lambert de Rouvroit C, Sluiters C, Cornelis GR. Role of the transcriptional activator, VirF, and temperature in the expression of the pYV plasmid genes of Yersinia enterocolitica. Mol Microbiol. 1992 Feb;6(3):395–409.
15. Jeckelmann JM, Erni B. Carbohydrate Transport by Group Translocation: The Bacterial Phosphoenolpyruvate: Sugar Phosphotransferase System. Vol. 92, Subcellular Biochemistry. Springer International Publishing; 2019. 223–274 p.
16. Jeckelmann JM, Erni B. Transporters of glucose and other carbohydrates in bacteria. Pflugers

Arch Eur J Physiol. 2020;472(9):1129–53.

17. Palace SG, Proulx MK, Lu S, Baker RE, Goguen JD. Genome-wide mutant fitness profiling identifies nutritional requirements for optimal growth of Yersinia pestis in deep tissue. MBio. 2014;5(4).
18. Wang Q, Millet YA, Chao MC, Sasabe J, Davis BM, Waldor MK. A genome-wide screen reveals that the Vibrio cholerae phosphoenolpyruvate phosphotransferase system modulates virulence gene expression. Infect Immun. 2015;83(9):3381–95.
19. Kok M, Bron G, Erni B, Mukhija S. Effect of enzyme I of the bacterial phosphoenolpyruvate : Sugar phosphotransferase system (PTS) on virulence in a murine model. Microbiology. 2003;149(9):2645–52.
20. Collins J, Robinson C, Danhof H, Knetsch CW, Van Leeuwen HC, Lawley TD, et al. Dietary trehalose enhances virulence of epidemic Clostridium difficile. Nature. 2018;553(7688):291–4.
21. Mazé A, Glatter T, Bumann D. The central metabolism regulator EIIAGlc switches salmonella from growth arrest to acute virulence through activation of virulence factor secretion. Cell Rep. 2014;7(5):1426–33.
22. Schuch R, Maurelli AT. The Mxi-Spa type III secretory pathway of Shigella flexneri requires an outer membrane lipoprotein, MxiM, for invasin translocation. Infect Immun. 1999;67(4):1982–91.
23. Burghout P, Beckers F, de Wit E, van Boxtel R, Cornelis GR, Tommassen J, et al. Role of the pilot protein YscW in the biogenesis of the YscC secretin in Yersinia enterocolitica. J Bacteriol. 2004 Aug 31;186(16):5366–75.
24. Rau R, Darwin AJ. Identification of YsaP, the pilotin of the Yersinia enterocolitica Ysa type III secretion system. J Bacteriol. 2015;197(17):2770–9.
25. Okon M, Moraes TF, Lario PI, Creagh AL, Haynes CA, Strynadka NC, et al. Structural Characterization of the Type-III Pilot-Secretin Complex from Shigella flexneri. Structure. 2008 Oct 8;16(10):1544–54.
26. Majewski DD, Okon M, Heinkel F, Robb CS, Vuckovic M, McIntosh LP, et al. Characterization of the Pilotin-Secretin Complex from the Salmonella enterica Type III Secretion System Using Hybrid Structural Methods. Structure. 2021 Feb;29(2):125-138.e5.
27. Carter PB, Zahorchak RJ, Brubaker RR. Plague virulence antigens from Yersinia enterocolitica. Infect Immun. 1980 May;28(2):638–40.
28. Sasakawa C, Kamata K, Sakai T, Murayama SY, Makino S, Yoshikawa M. Molecular alteration of the 140-megadalton plasmid associated with loss of virulence and Congo red binding activity in Shigella flexneri. Infect Immun. 1986 Feb;51(2):470–5.
29. Sturm A, Heinemann M, Arnoldini M, Benecke A, Ackermann M, Benz M, et al. The cost of virulence: retarded growth of Salmonella Typhimurium cells expressing type III secretion system 1. PLoS Pathog. 2011 Jul;7(7):e1002143.
30. Milne-Davies B, Helbig C, Wimmi S, Cheng DWC, Paczia N, Diepold A. Life After Secretion—Yersinia enterocolitica Rapidly Toggles Effector Secretion and Can Resume Cell Division in Response to Changing External Conditions. Front Microbiol. 2019 Sep 13;10:2128.

31. Kudryashev M, Diepold A, Amstutz M, Armitage JP, Stahlberg H, Cornelis GR. Yersinia enterocolitica type III secretion injectisomes form regularly spaced clusters, which incorporate new machines upon activation. Mol Microbiol. 2015 Dec 18;95(5):875–84.
32. Zhang Y, Lara-Tejero M, Bewersdorf J, Galán JE. Visualization and characterization of individual type III protein secretion machines in live bacteria. Proc Natl Acad Sci U S A. 2017 Jun 6;114(23):6098–103.
33. Gur E, Sauer RT. Recognition of misfolded proteins by Lon, a AAA+ protease. Genes Dev. 2008;22(16):2267–77.
34. Tsilibaris V, Maenhaut-Michel G, Van Melderen L. Biological roles of the Lon ATP-dependent protease. Res Microbiol. 2006;157(8):701–13.
35. Omnus DJ, Fink MJ, Szwedo K, Jonas K. The lon protease temporally restricts polar cell differentiation events during the caulobacter cell cycle. Elife. 2021;10:1–24.
36. Cornelis GR, Sluiters C, Delor I, Geib D, Kaniga K, de Rouvroit CL, et al. ymoA, a Yersinia enterocolitica chromosomal gene modulating the expression of virulence functions. Mol Microbiol. 1991 May;5(5):1023–34.
37. Cornelis GR. Role of the Transcription Activator VirF and the Histone-like Protein YmoA in the Thermoregulation of Virulence Functions in Yersiniae. Zentralblatt fur Bakteriol. 1993;278(2–3):149–64.
38. Jackson MW, Silva-Herzog E, Plano G V. The ATP-dependent ClpXP and Lon proteases regulate expression of the Yersinia pestis type III secretion system via regulated proteolysis of YmoA, a small histone-like protein. Mol Microbiol. 2004;54(5):1364–78.
39. Breidenstein EBM, Janot L, Strehmel J, Fernandez L, Taylor PK, Kukavica-Ibrulj I, et al. The Lon Protease Is Essential for Full Virulence in Pseudomonas aeruginosa. PLoS One. 2012;7(11).
40. Bretz J, Losada L, Lisboa K, Hutcheson SW. Lon protease functions as a negative regulator of type III protein secretion in Pseudomonas syringae. Mol Microbiol. 2002;45(2):397–409.
41. Zhou X, Teper D, Andrade MO, Zhang T, Chen S, Song WY, et al. A phosphorylation switch on lon protease regulates bacterial type iii secretion system in host. MBio. 2018;9(1).
42. Lee JH, Ancona V, Zhao Y. Lon protease modulates virulence traits in Erwinia amylovora by direct monitoring of major regulators and indirectly through the Rcs and Gac-Csr regulatory systems. Mol Plant Pathol. 2018;19(4):827–40.
43. Figaj D, Czaplewska P, Przepióra T, Ambroziak P, Potrykus M, Skorko-Glonek J. Lon protease is important for growth under stressful conditions and pathogenicity of the phytopathogen, bacterium Dickeya solani. Int J Mol Sci. 2020;21(10).
